# A dual-factor complex governs archaeal ribosome hibernation by sensing energy status

**DOI:** 10.64898/2026.01.19.700304

**Authors:** Lingfan Zhu, Rikuan Zheng, Keqin Wu, Xianshengjie Lang, Yueting Liang, Chuansen Yi, Qinqin Wang, Lei Qi, Jianyi Yang, Zhe Zhang, Xuben Hou, Jie Li, Chaomin Sun, Wenfei Li

## Abstract

Efficient coupling of cellular energy status to ribosome regulation is fundamental for cellular survival. Here, we identify the Archaeal Ribosome regulatory Complex (ARC)—a previously uncharacterized dual-factor complex comprising ARC-P and ARC-A. Cryo-EM reveals that ARC anchors to the small ribosomal subunit, establishing a stringent blockade of the tRNA and mRNA paths via an electrostatic wedge mechanism. This hibernation state is modulated by energy status: the complex remains stably associated under low-energy conditions, whereas ATP binding triggers rapid mobilization. Genetic analyses demonstrate that ARC acts as a translational brake essential for preserving ribosomal structural integrity, with ARC-A serving as the primary inhibitory factor. Notably, while the loss of ARC accelerates growth, it impairs recovery in proportion to stress severity. Phylogenomic analysis identifies ARC as an evolutionarily novel and widely distributed dual-factor system, revealing a conserved regulatory strategy for metabolic-translational coupling across diverse archaea. Our findings define a fundamental strategy where modular complexes integrate metabolic sensing with mechanical sequestration to balance growth and persistence.

## Introduction

Protein synthesis is one of the most energy-intensive processes in the cell. Synchronizing this expenditure with fluctuating energy availability is a fundamental requirement for fitness. Ribosomes, as the primary engines of translation and the major consumers of cellular energy, must be subject to stringent regulation to prevent catastrophic ATP depletion during environmental stress^1^. Under adverse environmental conditions, cells across Bacteria^2–9^, Archaea^10–12^, and Eukarya^13–18^ transition their ribosomes into a dormant state known as hibernation to minimize energy consumption^19–23^. Despite the universality of this strategy^24–27^, the mechanisms transducing energy status into translational arrest remain largely unexplored.

In archaea, a unique evolutionary lineage that bridges bacterial simplicity and eukaryotic complexity^28–31^, the mechanisms of ribosome hibernation are still largely unexplored. Recent studies, focused primarily on thermophilic archaea, identified single-protein factors as key mediators of ribosomal dormancy^10–12^ through mechanisms such as ribosome dimerization or anti-association^29,32^. However, the regulatory diversity across the vast archaeal landscape, especially for organisms inhabiting extreme and fluctuating environments like deep-sea cold seeps, remains to be fully uncovered. Methanogens, in particular, constitute a globally distributed phylum adapted to strictly anaerobic and chronically energy-limited environments^31,33–35^, where any delay in metabolic adaptation during starvation can be lethal. Among these, the deep-sea methanogen *Methanolobus* sp. ZRKC1 (hereafter referred to as *Methanolobus*), isolated from a newly formed cold-seep vent field, presents a unique metabolic profile utilizing complex polysaccharides that necessitate high-cost enzyme synthesis^36^ (Fig. 1A). Given the unpredictable and often delayed supply of key metabolites such as methanol in dynamic cold-seep environments, we reasoned that this organism must have evolved a robust dormancy mechanism to instantaneously suppress translation and preserve essential high-energy reserves, such as polyphosphate, against wasteful depletion during periods of starvation^36^.

**Fig. 1.**
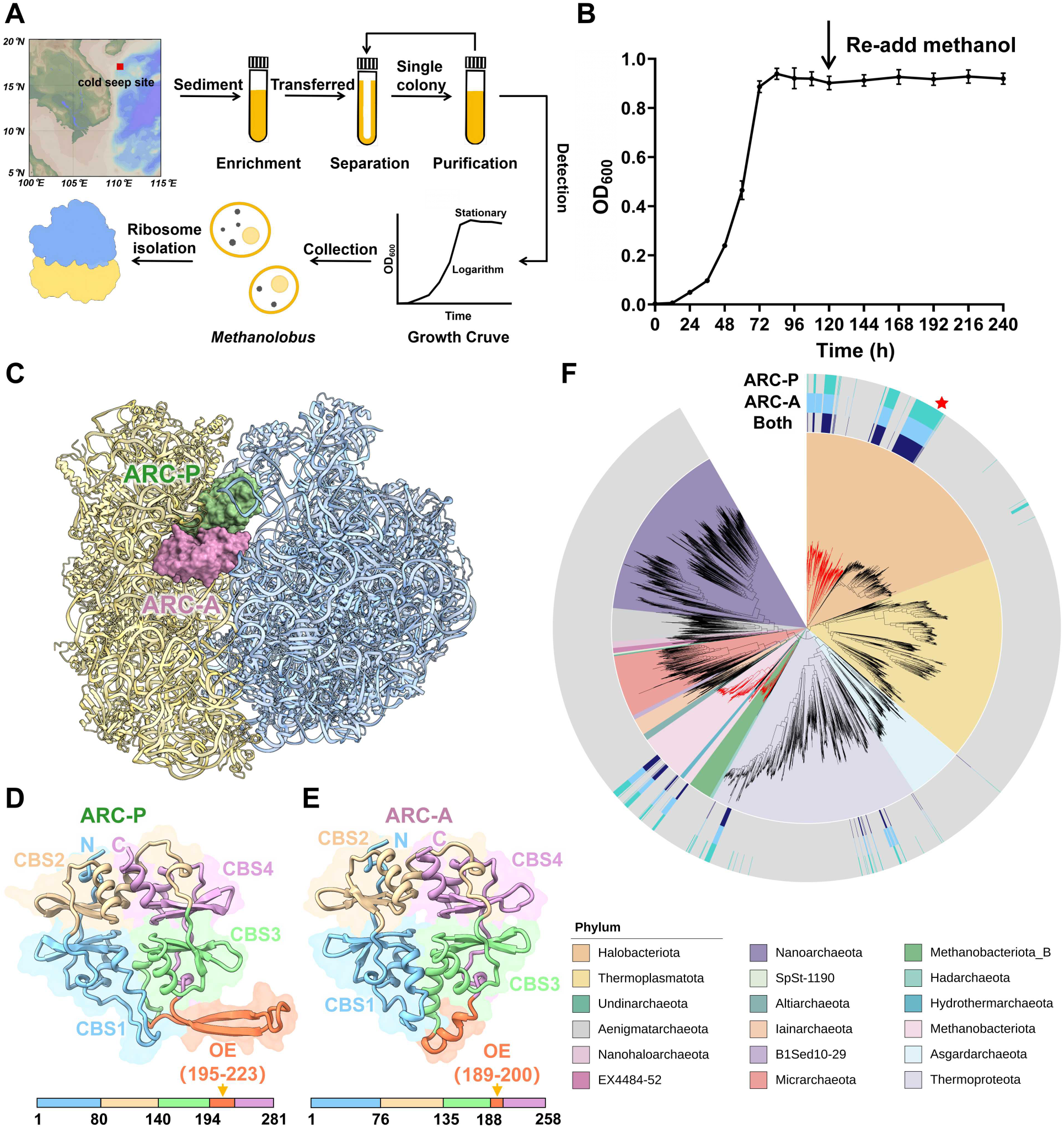
Discovery, structural organization, and phylogenetic analysis of the ARC. (**A**) Schematic workflow of *Methanolobus* isolation and sample preparasion for cryo-EM analysis.This map was generated using Ocean Data View^39^. (**B**) Growth profile of *Methanolobus* with methanol re-supplementation during stationary phase. Methanol was re-added at the indicated time point (black arrow). (**C**) Cryo-EM structure of ARC bound to the non-rotated *Methanolobus* 70S ribosome with ARC-A (pink) and ARC-P (green) shown in surface, and the ribosome in cartoon. (**D and E**) Cryo-EM structures and schematic domain organization of ARC-P (D) and ARC-A (E) with the corresponding sequence regions indicated in the bar diagram. OE, occlusion element. (**F**) Phylogenetic distribution of ARC-P and ARC-A homologs across the Archaea domain. The outer, middle, and inner rings denote organisms possessing homologs of ARC-P, ARC-A, and both, respectively. Most methanogenic archaea are represented by red lines, and *Methanolobus* is indicated by a red star. Note that, methanogens are distributed across multiple phyla (such as *Halobacteriota* and *Methanobacteriota*)^40^.

Intriguingly, we observed that *Methanolobus* cells entering the stationary phase exhibit a characteristic growth inertia: supplementation with fresh metabolic substrates fails to restore biomass accumulation (Fig. 1B). This persistent lag suggests that the translational machinery is not merely stalled by substrate depletion but is actively sequestered by a potent hibernation system to prioritize long-term persistence over opportunistic growth. To elucidate the mechanisms of this arrest, we characterized the ribosomal assemblies of *Methanolobus* as it transitioned from active growth to maintenance mode. This led to the identification of ARC (Archaeal Ribosome regulatory Complex), a unique dual-factor system comprising ARC-P and ARC-A. Our findings define ARC as a distinctive module that enforces stringent translational arrest, revealing a previously uncharacterized strategy for ribosome hibernation that couples ancient metabolic sensing with the universal machinery of protein synthesis.

## Results

### Discovery of the dual-factor ARC in dormant ribosomes

To define the metabolic transition triggering ribosomal arrest, we first characterized the growth dynamics of *Methanolobus*. When grown on methanol, cells reached a stable stationary phase upon substrate exhaustion, transitioning into a maintenance mode sustained by the catabolism of internally stored polyphosphate (Fig. 1A and 1B, fig. S1A). Notably, the late stationary-phase cells exhibited significant growth inertia, where re-supplementation with methanol failed to trigger immediate biomass accumulation (Fig. 1B), suggesting a sequestered translation machinery. Sucrose gradient analysis revealed a marked shift in the ribosomal profile during the transition to dormancy (fig. S1B). Logarithmic cells exhibited a prominent pool of free small ribosomal subunits (SSU) relative to the combined peak of large subunits (LSU) and 70S monosomes. In contrast, stationary-phase extracts showed a significant reduction in the SSU proportion, indicating the systematic sequestration of ribosomal components into stable 70S assemblies.

To understand the molecular basis of this state, we performed cryo-EM analysis of ribosomes extracted from this late stationary-phase cells. Subsequent data analysis yielded a complete 70S ribosome structure at 2.3 Å resolution, with local resolution reaching up to 2 Å (figs. S1 and S2, tables S1 and S2), enabling accurate *de novo* atomic modeling of the three rRNAs and 56 annotated ribosomal proteins (fig. S3A). The high-quality map further allowed us to resolve archaea-specific components, including aS21^10,37^, a novel SSU protein aS22 (DUF5350), and ten well-defined, coordinated zinc ions (fig. S3B to D).

The dominant 70S class (84.3% of the 70S monomer) revealed two prominent densities at the subunit interface (Fig. 1C). Sequence alignment and subsequent modeling unambiguously identified these as a novel, two-component complex of Cystathionine Beta-Synthase (CBS) domain-containing proteins^38^, which we term the ARC (Archaeal Ribosome regulatory Complex) (Fig. 1D and E). The ARC complex is composed of two non-fused, previously unannotated multi-domain CBS proteins: ARC-P and ARC-A. Each compoent contains four CBS domains and a unique occlusion element (OE) comprising compact β-sheets and α-helices, respectively (Fig. 1D and E). Structural analysis reveals that ARC-P and ARC-A exhibit high structural homology (RMSD of 1.2 Å) (fig. S3E). This architecture represents a fundamental divergence from the single-polypeptide hibernation models previously characterized in Archaea^10,11^.

Phylogenetic analysis reveals that while ARC-P and ARC-A share the conserved CBS domain structure, a module widely recognized for binding nucleotides and coupling cellular energy status to protein function across domains of life, they represent a distinct evolutionary lineage from previously discovered archaeal single-polypeptide ribosome hibernation factors^10,11^. ARC-P exhibits a remarkably broad distribution across diverse archaea phyla, including *Methanobacteriota*, *Thermoproteota* and *Halobacteriota*, as well as specific bacteria (fig. S4A, data S1 and S2). ARC-A is also present in multiple archaeal and bacterial phyla, yet, it forms a unique and novel evolutionary clade, supporting its designation as a newly discovered family (fig. S4B, data S3 and S4). Crucially, the genomic co-occurrence of *arcP* and *arcA* is a highly conserved and dominant feature across major archaeal phyla, supporting their co-evolution as a single functional, cooperative module (Fig. 1F, data S5).

### Cooperative stabilization and molecular basis for ARC anchoring

To determine how ARC engages the translational machinery across physiological states, we monitored its occupancy across different growth stages, ranging from early logarithmic phase to deep stationary phase (Fig. 2A and B). Immunoblotting of sucrose gradient fractions at various growth stages (OD_600_ 0.2, 0.4, 0.6 and 0.9) revealed that both ARC-A and ARC-P remain constitutively associated with the SSU even during active growth. Interestingly, while ARC remains bound to the SSU pool throughout the logarithmic phase, its occupancy in the total ribosome fraction significantly enriches as cells transition from active growth into the maintenance mode of the stationary phase (Fig. 2B). This constitutive association identifies ARC as a pre-positioned sentinel rather than a factor recruited *de novo* upon energy depletion, ensuring an instantaneous translational response to fluctuating environmental conditions.

**Fig. 2.**
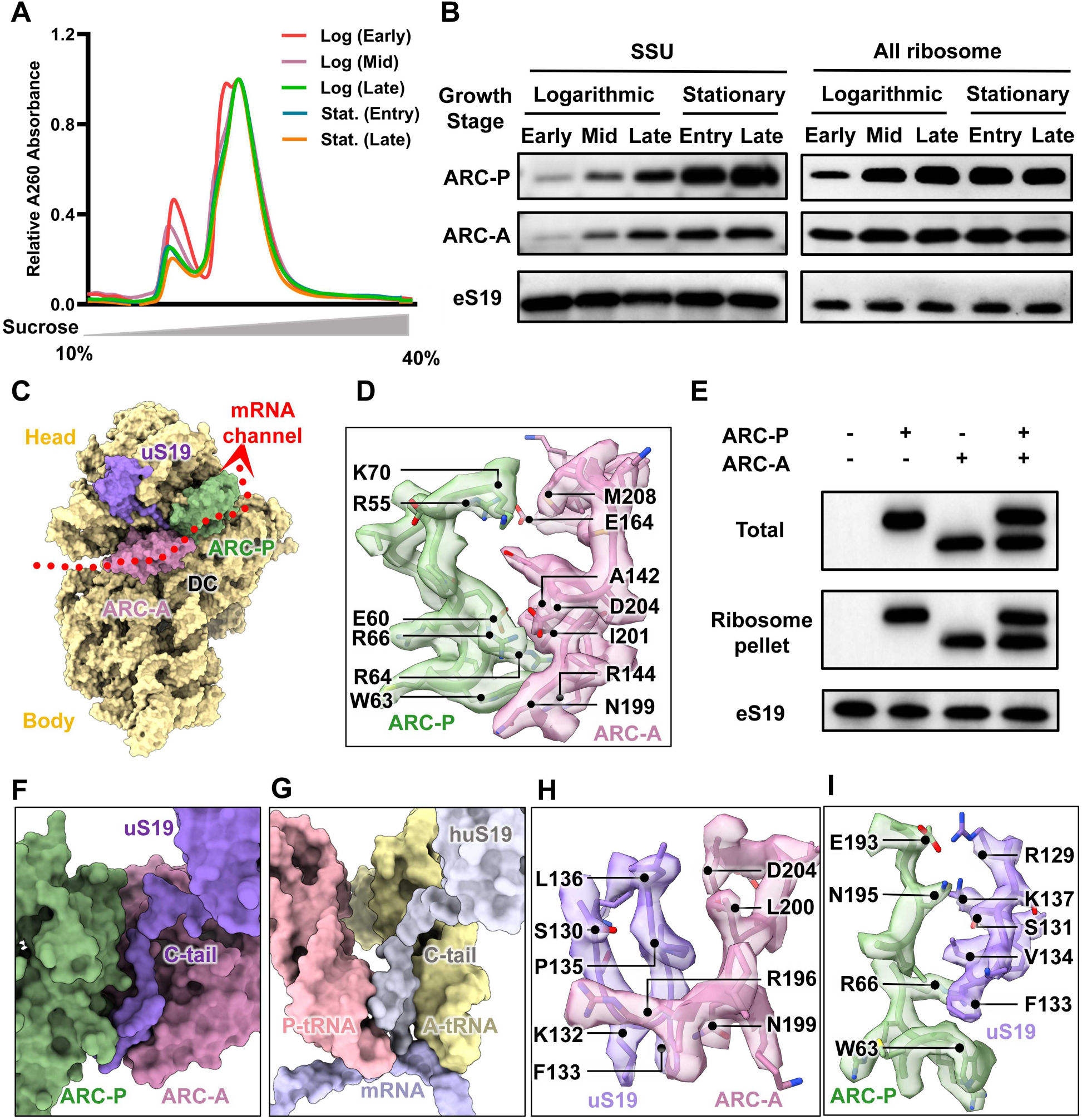
Molecular basis of constitutive ARC anchoring and cooperative recruitment. (**A**) Representive sucrose gradient profile (10%-40%) of cell extracts across different growth stages: early, mid, and late logarithmic phases (OD_600_ = 0.2, 0.4, 0.6, respectively) as well as entry/late stationary phases. Log, logarithmic phase. Stat, stationary phase. (**B**) Western blot analysis and quantification of ARC occupancy on the SSU and total ribosome fractions across growth phases. (**C**) Position of ARC-P and ARC-A on the small ribosomal subunit (SSU). All components are shown in surface. The path of the mRNA channel path is indicated by a red dashed line. DC, Decoding Center. (**D**) Close-up view of the interface between ARC-A and ARC-P. (**E**) Ribosome binding assays showing the co-sedimentation of ARC proteins with the ribosome. ARC proteins are FLAG tagged and visualized using an anti-FLAG antibody. (**F**) Association of the ARC complex with the uS19 C-tail in the non-rotated 70S ribosome. (**G**) Comparisons of the uS19 C-tail position in the human ribosome (PDB: 6Y0G^42^), where it is inserted between the tRNA and mRNA. (**H and I**) Key molecular interactions between uS19 and ARC-A (H) or ARC-P (I). Proteins are shown in cartoon representation with corresponding cryo-EM density.

To investigate the physical basis for this dual-factor regulation, we employed CryoDRGN analysis^41^, which confirmed that the ARC complex predominantly exists as a stable, cooperative bound assembly on the 70S ribosome (fig. S1E). The ARC factors occupy functionally distinct sites (Fig. 2C): ARC-P acts as the exit pathway gatekeeper near the P and E-sites, while ARC-A specifically engages the A-site (Fig. 2C, fig. S5A). The stability of this dual blockade relies on extensive cooperative protein-protein interactions (Fig. 2D). Ribosome binding assays further confirmed that while both ARC-P and ARC-A could bind to the ribosome individually *in vitro*, co-addition resulted in co-sedimentation of both factors on the ribosome (Fig. 2E).

Anchoring of ARC is facilitated by the ribosomal protein uS19, whose extended C-terminus tail (C-tail) directly serves as a molecular tether to secure the ARC complex in its arrested conformation (Fig. 2F). This interaction remarkably mirrors the role of the uS19 C-tail in stabilizing tRNA accommodation within mammalian ribosomes^42^ (Fig. 2G), with the interface further stabilized by specific molecular forces including a key hydrogen bond between N195 (ARC-P) and K137 (uS19) (Fig. 2H and I). The conserved role of uS19 in stabilizing functionally distinct ribosomal conformations across archaea and eukaryotes suggests a deep evolutionary origin for this regulatory hub (fig. S5B and C).

### ARC enforces dual blockade of the mRNA channel and critical A-site remodeling

The dual occupancy of ARC establishes a comprehensive two-point blockade of the ribosome’s functional centers, effectively sealing the A-, P-, and E-sites alongside the entire mRNA channel (Fig. 3A). This stringent inhibition is achieved through the rigid occlusion elements (OEs), which extend deeply into the mRNA channel. We found that the stability of this blockade relies on a remarkable electrostatic complementarity. The highly basic OEs (harboring concentrated KRK clusters) project into the polyanionic environment of the rRNA, effectively functioning as an electrostatic wedge that anchors the complex within the decoding center (fig. S6).

**Fig. 3.**
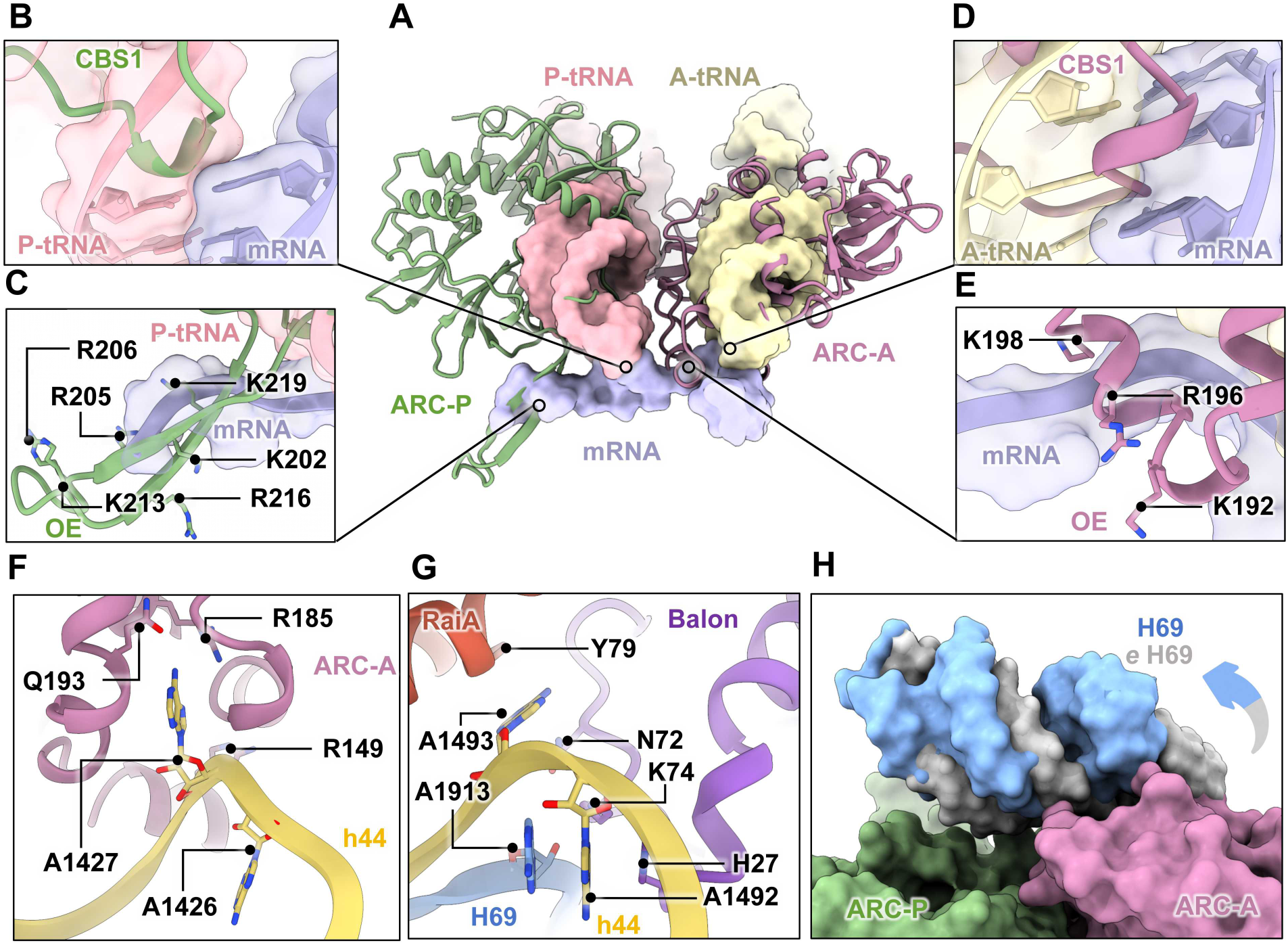
Molecular mechanism of dual blockade and decoding center rearrangement by ARC. (**A**) Superimpose of ARC positions on the 70S ribosome relative to the tRNA binding sites (PDB: 7K00^14^). (**B and C**) A zoom-in view of ARC-P interfering with the P-tRNA/mRNA pairs via its CBS1 domain (B), and sealing the mRNA channel via its OE (C). (**D and E**) Close-up views of ARC-A disrupting the A-tRNA/mRNA interaction via its CBS1 domain (D) and inserting its OE into the mRNA channel (E). Basic residues (K/R) serving as electrostatic anchors are indicated by sticks in (C) and (E). (**F**) Induced outward flip of the A1427 and inward flip of A1426 of h44 upon ARC-A binding. (**G**) Structural comparison with the bacterial hibernation factor Balon (PDB: 8RD8^5^), showing convergent remodeling of the monitoring bases (A1493/A1492 of h44). The A1913 of H69 and the A1492 of h44 form π-π stacking. (**H**) Retraction of the inter-subunit bridge H69 toward the LSU relative to the bacterial ribosome (PDB: 7K00^14^) to accommodate the ARC-A. OE, occlusion element.

ARC-P executes the exit-site blockade, obstructing the P-tRNA path via its CBS domains and the mRNA exit channel through its OE (Fig. 3B and C). This element is flanked by pairs of Lysine (K) and Arginine (R) residues serving as an electrostatic anchor within the narrowest constriction of the mRNA exit tunnel (Fig. 3C, fig. S6A to B). Simultaneously, ARC-A occupies the A-tRNA binding pocket through its CBS1 domain, sterically precluding tRNA association (Fig. 3D and E). Its OE—harboring a concentrated basic cluster (KRK)—inserts directly into the decoding center and the mRNA channel (Fig. 3E, fig. S6A and C). This electrostatic engagement provides a second robust anchor that induces a critical molecular remodeling of the decoding center. Specifically, The binding of ARC-A induces a dramatic outward conformational flip of the A1427 residue (corresponding to the bacterial A1493) in helix 44 (h44) (Fig. 3F, fig. S6D). This rearrangement is a hallmark of stringent translational arrest abserved across domains ^5,43,44^. Furthermore, the inter-subunit bridge H69 undergoes a significant conformational rearrangement, retracting towards the LSU to avoid a steric clash with the deeply inserted ARC-A (Fig. 3H).

Notably, structural comparisons confirmed that ARC-A sterically mimics both canonical A-site substrates (such as tRNA and eRF1^45^) and A-site hibernation factors like Balon^5^. Despite their distinct evolutionary origins, both systems converge on the induction of the critical outward flip of the monitoring base (A1427/A1493) to block decoding (Fig. 3F to G, fig. S7), suggesting a universally conserved requirement for translational arrest.

Together, these structural observations characterize ARC as a dual-action inhibitor. It effectively functions as a physical plug that seals the mRNA path while simultaneously acting as a molecular switch that deactivates the decoding mechanism through specific rRNA remodeling. This dual strategy provides a molecular basis for the robust growth inertia observed in our physiological assays, as the resumption of protein synthesis would require the synchronized removal of both the physical and electrostatic seals.

### ARC functions as a metabolic sensor coupling energy status to translation

The conserved CBS domains suggested ARC acts as a nucleotide-dependent metabolic switch^11,46^. To define the molecular basis for this energy sensing, we determined the cryo-EM structure of the ARC-bound 70S ribosome in the presence of AMP (fig. S8, tables S1 and S2). The map clearly resolved three AMP molecules sequestered within the CBS pockets: two in ARC-P (CBS1 and CBS2) and one in ARC-A (CBS3) (Fig. 4A). This coordination mirrors the conserved logic of eukaryotic energy sensors like the γ subunit of AMPK^47^ where nucleotide binding is mediated by conserved polar residue pairs like Threonine-Aspartate (T/D) pairs (Fig. 4B, fig. S9). In ARC-A, D176 in CBS3 provides a critical anchor for AMP, while an Asp-to-Asn substitution (N50) explains the absence of nucleotide density in its CBS1 (Fig. 4B, fig. S9A and B). AMP within ARC-A is stabilized by sophisticated coordination network featuring prominent π-stacking interaction, contrasting with the simpler hydrogen-bonding networks utilized by ARC-P (fig. S9B to D). Notably, AMP binding does not induce significant global conformational shifts, suggesting that the hibernation complex maintains a rigid and stable architecture under low-energy conditions.

**Fig. 4.**
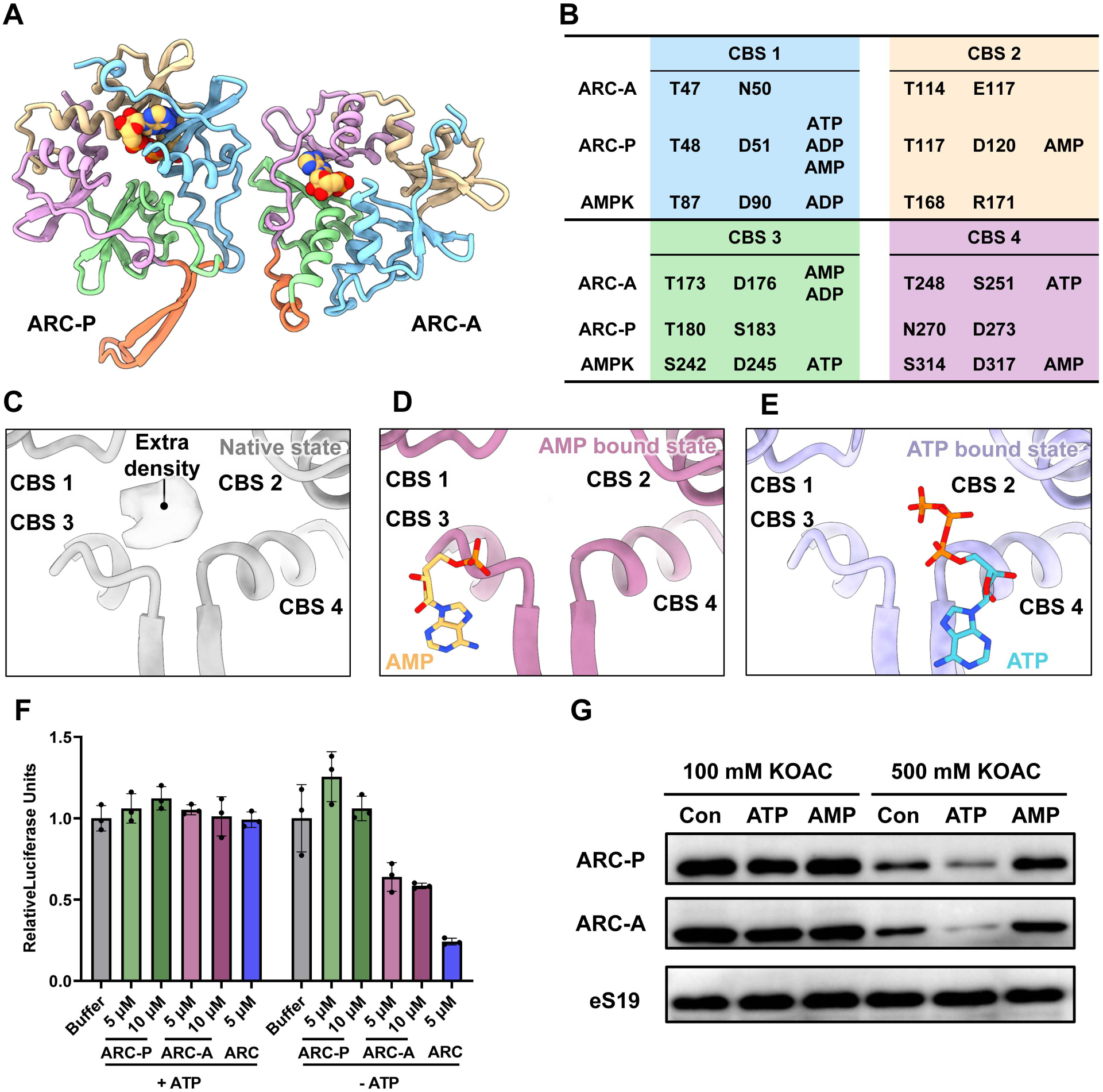
Nucleotide-dependent regulation and energy sensing by ARC. (**A**) Atomic model of the AMP-bound ARC. (**B**) Table of conserved amino acid residues critical for AMP coordination within the binding pockets of ARC and the eukaryotic AMPK γ subunit. (**C to E**) Zoom-in views of the ARC-A arginine pocket in native (C), AMP bound (D) and ATP bound (E, predicted by AlphaFold 3^48^) states. (**F**) IVT assays showing the nucleotide-dependent inhibition of Nano-luciferase reporter efficiency in the *Methanolobus* cell-free system. Data are presented as mean ± s.d. (n=3). (**G**) Western blot analysis of ARC–ribosome binding stability using sucrose cushions at increased salt concentrations in the presence of 2 mM ATP or AMP, illustrating the nucleotide-specific mobilization of the complex from the 70S ribosome.

Comparison between the native and AMP-reconstituted structures reveals a regulatory hub unique to ARC-A. The native 70S-ARC complex exhibits an unexplained density within a positively charged arginine pocket (Fig. 4C, fig. S10A). This density is absent in the AMP-reconstituted system despite the saturation of adjacent nucleotide-binding sites (Fig. 4D, fig. S10B). Structural modeling demonstrates that ATP binds at the CBS4 site, distinct from the stable AMP-binding pockets (Fig. 4E, fig. S10C), where its triphosphate group precisely occupies the volume of this native density (fig. S10D to E). Furthermore, the concentrated positive charge of this arginine cluster provides a highly complementary environment for the negatively charged phosphate tail of ATP (fig. S10D to E). The specific vacancy of this pocket under high AMP concentrations identifies it as a high-threshold sensing gate in a primed state, enabling the regulatory transition upon ATP recognition.

The functional consequences of this energy sensing were evaluated through *in vitro* translation (IVT) assays (Fig. 4F). In the presence of ATP, the ribosome remains insensitive to ARC-mediated arrest, reflecting active-state stabilization where high energy charge prevents inhibitory binding (Fig. 4F). Conversely, ATP depletion reveals a different regulatory profile in which ARC-P alone exhibits negligible inhibitor capacity whereas ARC-A independently supressing translation by approximately 50%. Their co-addition elicits a potent synergistic arrest that cooperatively locks the ribosome (Fig. 4F), ensuring complete translational shutdown during energy scarcity. To confirm that this recovery stems from physical dissociation, we performed biochemical mobilization assays under varying ionic strengths. While ARC remained anchored with AMP, ATP triggered a significant, salt-sensitive release of both subunits into the supernatant (Fig. 4G). This ATP-specific destabilization of the electrostatic anchor identifies ARC as a reversible, energy-sensitive brake that is dismissed upon metabolic recovery. Molecular dynamics (MD) simulations further revealed that while AMP occupancy stabilizes the complex, ATP binding at the CBS4 sensing site significantly increases the root-mean-square fluctuation (RMSF) of the OEs (fig. S10G and H). This binding event neutralizes the arginine pocket, providing the allosteric impetus to mobilize the electrostatic wedge and facilitate the rapid resumption of protein synthesis (fig. S10F).

### ARC-mediated ribosome sequestration safeguards structural integrity and fitness during metabolic transitions

To investigate the physiological consequences of ARC, we established the first successful genetic manipulation of this environmentally critical, deep-sea methanogen (Fig. 5A, fig. S11A to C). We generated single and double deletion strains for *arcP* and *arcA* to examine their impact on cellular fitness. Surprisingly, under nutrient-replete conditions during the exponential phase, genetic ablation of the ARC complex significantly accelerated biomass accumulation compared to the wild-type (WT) (Fig. 5B). A clear funcational hierarchy was observed among the mutants: the double-deletion strain (Δ*arc*) exhibited the most rapid growth, followed by Δ*arcA*, while Δ*arcP* showed a more modest growth increase. These data, coupled with our observation that ARC remains associated with the SSU during active growth (Fig. 2B), demonstrate that ARC functions as a constitutive translational brake that imposes a deliberate metabolic drag on the cell.

**Fig. 5.**
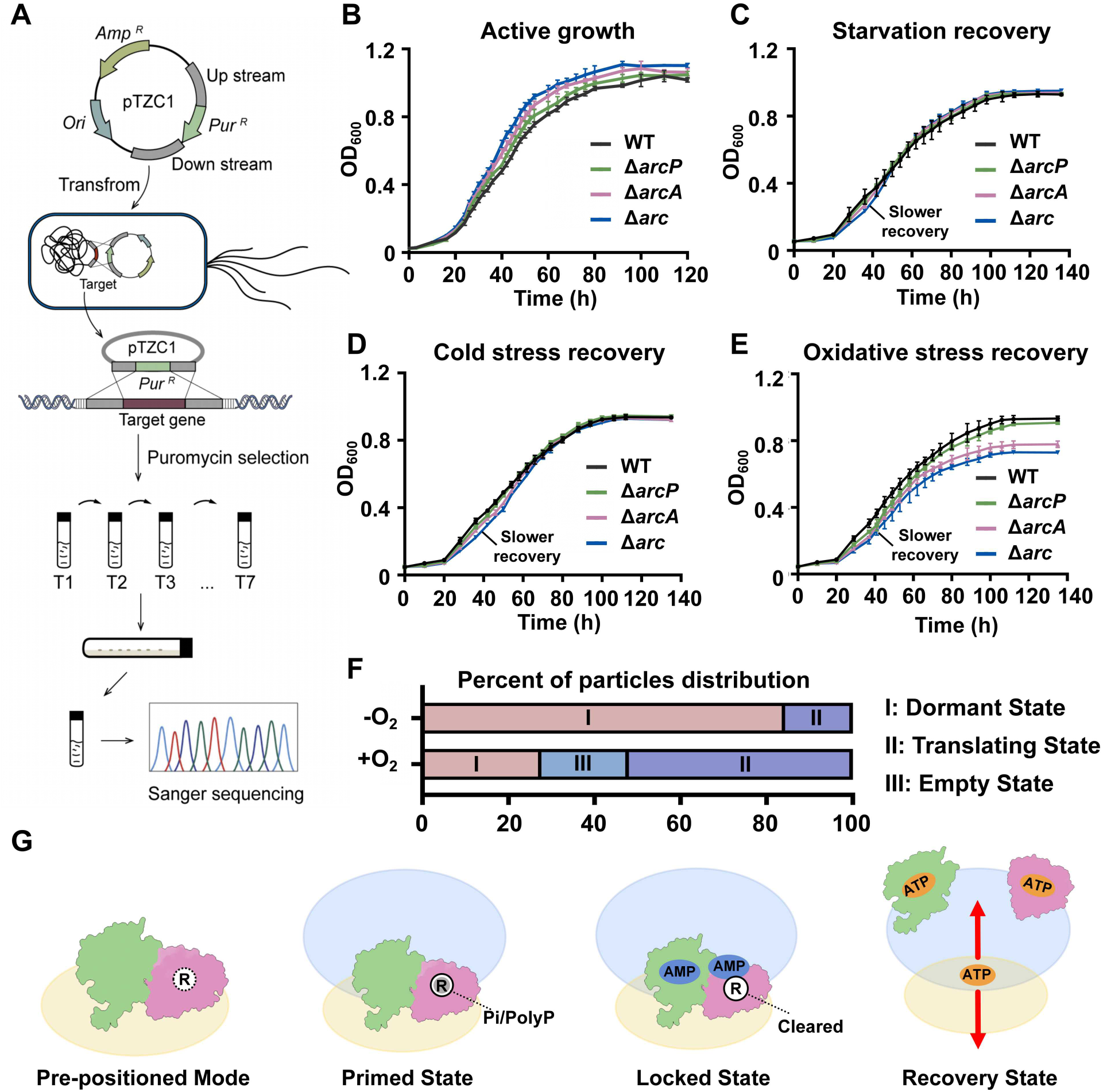
ARC-mediated translational arrest and metabolic mobilization. (**A**) Schematic of the genetic manipulation system developed for *Methanolobus*. (**B to E**) Growth profiles of the wild-type (WT) and ARC-deficient strains (Δ*arc,* Δ*arcA* and Δ*arcP*) inoculated from different states: mid-logarithmic phase (B), stationary-phase cells (C), cold stress (D), and oxygen stress (E). (**F**) Quantitative distribution of 70S ribosomal classes during the transition from dormancy to mobilization. (**G**) Proposed schematic model of ARC-mediated metabolic-translational switch. The model illustrates the occupancy of CBS domains by AMP during energy scarcity and the ATP-driven mobilization during metabolic recovery.

However, this opportunistic growth acceleration came at a profound cost to environmental resilience. When transitioning from prolonged dormancy in the stationary phase or encountering cold stress, the ARC-deficient strains consistently exhibited a pronounced recovery lag compared to the WT, although they eventually attained a similar final biomass (Fig. 5C and D, fig. S11D and E). Across these scenarios, the severity of the recovery deficit mirrored the inhibitory potency of the factors, identifying ARC-A as the primary determinant of post-stress fitness. Strikingly, under acute oxygen stress, a condition particularly detrimental to these anaerobic methanogens, the growth hierarchy observed during active proliferation was completely reversed (Fig. 5B and E). While the WT achieved the most rapid regrowth and the highest final biomass, the ARC-deficient mutants displayed an extensive recovery lag and signicantly reduced final biomass (Fig. 5E).

To determine how metabolic disruption influences the stability of the ARC-ribosome complex, we utilized the acute physiological response to oxygen exposure—which induced the most severe metabolic remodeling in our assays—to capture cryo-EM snapshots of ribosomes transitioning toward mobilization (fig. S12, table S3). Quantitative classification revealed a marked shift in the ribosomal landscape: ARC occupancy dropped significantly, accompanied by a synchronized increase in translating 70S ribosomes (Fig. 5F). Crucially, 3D variability analysis (3DVA)^49^ revealed that ribosomes lacking ARC occupancy in these mobilizing cells exhibit pronounced conformational flux and structural disorder within the SSU head region (movie S1).

This structural vulnerability provides a physical explanation for the physiological failure of the Δ*arc* mutants. Without the mechanical support provided by the electrostatic wedge of ARC, the SSU head becomes prone to irreversible structural disarray during energy-limited transitions. These findings identify ARC not merely as a passive inhibitor, but as a structural guarantor that couples metabolic sensing to ribosomal preservation. By enforcing a protected, 70S-sequestered state, ARC prevents the catastrophic degradation of the translational machinery, ensuring that the cell remains primed for rapid and synchronized recovery upon environmental restoration (Fig. 5G).

## Discussion

The discovery of ARC reveals a sophisticated dual-factor providing the critical molecular link that allows the ribosome to sense and respond to cellular energy status. While previously characterized hibernation factors, such as Hib and Dri^10,11^, typically rely on single-polypeptide architectures to enforce growth arrest, ARC utilizes a cooperative assembly of distinct subunits to achieve a comprehensive blockade of the ribosome, ensuring that the translational machinery remains protected during the prolonged nutrient limitation.

Our phylogenomic analysis underscores the significance and broad distribution of this discovery. We identified ARC components across diverse archaeal phyla, revealing a striking mutual exclusivity between the ARC-A sensor module and the previously characterized Dri protein^10^. The observation that *dri* occurrence drops to zero in lineages harboring *arcA* identifies this dual-factor system as a specialized functional successor, or evolutionary upgrade, optimized for energy-limited ecosystems (data. S3 and S4). This distribution suggests that ARC is not an isolated case but a fundamental regulatory system that has likely rendered ancestral, single-factor mechanisms redundant in deep-sea environments.

The core logic of the ARC system rests on the coupling of energy-sensing to structural sequestration. Rather than a stress-induced recruitment, the ARC complex maintains a constitutive presence on the small ribosomal subunit across all growth phases, ensuring the translational machinery is always primed for an instantaneous response to energy fluctuations. Under low-energy conditions, the stable association of ARC rigidifies the complexthrough its electrostatic wedge mechanism. Upon the restoration of energy levels, the binding of the negatively charged triphosphate moiety of ATP to the sensor pocket of ARC-A neutralizes the anchoring forces, providing the allosteric impetus for mobilization (fig. S10G and H). This coupling, integrated into our comprehensive model of energy-dependent ribosome sequestration (Fig. 5G), ensures that the ribosome remains unresponsive to potential substrates only as long as the energy charge remains below a critical threshold.

This regulatory coupling also underscores a fundamental principle of energy management, wherein structural preservation is prioritized over immediate proliferation. Our genetic data indicate that growth inertia in *Methanolobus* represents a regulated state of persistence — an evolutionary trade-off that favors long-term viability over opportunistic growth. By sequestering ribosomes in a protected assembly, ARC prevents structural degradation of the translational apparatus during starvation. The constitutive presence of ARC results in a reduction in the maximal growth rate during proliferation.

Ultimately, the integration of this ancient metabolic sensing module into the translation apparatus suggests that the coupling of energy status to structural integrity is a fundamental evolutionary adaptation, representing a conserved mechanism for life under extreme thermodynamic constraints.

## Methods

### *Methanolobus* isolation, culture and stress treatment

To isolate and cultivate deep-sea methanogenic archaea, the cold seep sedimental samples (E110°35’02.215”, N17°07’15.236”) were added to the 15 mL anaerobic tube containing 10 mL basal medium (20.0 g/L NaCl, 1.0 g/L yeast extract, 1.0 g/L peptone, 1.0 g/L NH_4_Cl, 1.0 g/L CH_3_COONa, 1.0 g/L NaHCO_3_, 0.2 g/L MgSO_4_, 0.1 g/L CaCl_2_, 1.0 g/L cysteine hydrochloride, 500 µL/L 0.1 % (w/v) resazurin, 1.0 L distilled water, pH 7.0) supplemented with 250 mM methanol and 100 µg/mL rifampicin. The inoculated media were incubated anaerobically at 28 °C for one month. The deep-sea methanogenic Archaea were identified by the polymerase chain reaction (PCR). Purity was confirmed regularly via transmission electron microscope (TEM) and 16S rRNA gene sequencing. Through isolation and identification, a novel methanogenic archaeal strain was obtained and named *Methanolobus* sp. ZRKC1 (GenBank: CP146978).

To establish sufficient biomass for downstream analyses, the isolated strain was expanded anaerobically at 28 °C in a defined basal medium, with distilled water replaced by filtered seawater (pH 7.0) to better mimic its natural habitat. Pre-cultures (4 mL) were used to inoculate 500 mL Hungate bottles containing 400 mL medium supplemented with 100 mM methanol. Cultures were incubated anaerobically at 28 °C without shaking. Growth was monitored by measuring OD_600_ values using a microplate reader (Infinite M1000 Pro; Tecan, Mannedorf, Switzerland). Methanol was re-added at the indicated time point. Upon the OD_600_ value reaching 0.9 (stationary phase), cultures designated for the non-stress (stationary phase) condition were immediately harvested by centrifugation at 4,000 *g* for 10 min at 4 °C.

For the oxidative stress treatment, anaerobic bottles were exposed to ambient air and agitated every 10 min to ensure oxygen saturation. Oxygen dissolution was verified by the medium’s colour change from colorless to pink (resazurin serving as the oxygen indicator), and the cultures were then incubated for 1 h before harvesting.

### *Methanolobus* ribosome isolation and purification

To isolate the *Methanolobus* ribosomes, the cell pellet was resuspended in lysis buffer (20 mM HEPES-KOH pH 7.5, 10 mM Mg(OAc)_2_, 100 mM NH₄Cl, 1 mM DTT). Cells were lysed using a high-pressure homogenizer (1290 bar, 4 passes), followed by centrifugation at 20,000 *g* for 10 min at 4 °C to remove cellular debris as described previously with minor modifications^50^. The supernatant was then loaded onto a 0.8 M sucrose cushion and centrifuged using an MLA130 rotor at 603,000 *g* for 50 min at 4 °C. The pellet was resuspended in 500 μL lysis buffer, and the crude ribosome sample was applied to a 10–40% sucrose density gradient (prepared in lysis buffer). The gradient was centrifuged using a Beckman-Coulter SW41 rotor at 247,600 *g* for 2.5 h. Fractions were analyzed by 2% agarose gel electrophoresis to verify the distribution of ribosomal species. Fractions corresponding to the 70S was pooled, pelleted through a 0.8 M sucrose cushion (MLA130 rotor, 603,000 *g*, 50 min, 4 °C) and the resulting pellet was resuspended in grid buffer (20 mM HEPES-KOH pH 7.5, 10 mM Mg(OAc)_2_, 30 mM KCl, 1 mM DTT) for cryo-EM grid freezing.

### Cryo-EM sample preparation and data acquisition

The concentrations of the *Methanolobus* ribosomes used for cryo-EM grid preparation were adjusted to A260 ≈ 8. Approximately, 3.5 μL aliquots were applied to a plasma-cleaned holey carbon grid (R1.2/1.3, 300 mesh, Quantfoil) for 1 min, on which a homemade continuous carbon film was precoated. Grids were blotted for 2.5 s under 100% humidity at 4 °C and plunge-frozen in liquid ethane using Vitrobot Mark IV (Thermo Fisher Scientific).

Data acquisition was performed on a Titan Krios transmission electron microscope (Thermo Fisher Scientific) operating at 300 kV, equipped with a GIF quantum energy filter (Gatan) and a 20 eV^-2^ slit width. Micrographs were captured utilizing a Falcon 4i direct electron detector, resulting in a physical pixel size of 0.93 Å. The total electron exposure was set at 60 electrons per Å^2^, distributed across 40 fractional frames over a 5.9 s total exposure time. Automated data collection utilized EPU software^51^, with defocus values spanning 1.0 to 2.0 μm.

### Cryo-EM data processing and structural analysis

The datasets were processed starting in cryoSPARC v.4.6.2^49^. Initial steps involved correcting beam-induced motion via patch motion correction and estimating CTF parameters using the patch CTF estimation tool. Micrographs were manually inspected and filtered for quality. Particle extraction commenced with template-free blob picking, refined by iterative template-based picking from averaged 2D class projections. Subsequent extensive 2D classification rounds were used to eliminate contamination and ice artifacts. The cleaned particle set was then subjected to initial 3D auto-refinement to generate a consensus map. For subsequent particle sorting, focused 3D classification with specific masks were employed. To enhance local resolution at key functional sites, spherical masks were generated using the “relion_mask_create” tool in RELION 4.0.1^52^ and applied during local focused refinement of each class. Following CTF refinement, particles from each resulting class underwent final masked refinements. Map resolutions are reported based on the FSC = 0.143 criterion.

### CryoDRGN conformational heterogeneity analysis

To analyze particles dynamics and heterogeneity, the consensus refined particles were subjected to CryoDRGN analysis^41^. The particles were down-sampled to a box size of 256 pixels, and twenty-five iterative rounds of the CryoDRGN VAE were trained using a network. The resulting z values were classified using k-means clustering and Gaussian mixture models and plotted using a UMAP approach.

### Model building and refinement

Genome sequencing data from *Methanolobus* were used to obtain rRNA and protein sequences. rRNA secondary structure was predicted using the R2DT tool (https://rnacentral.org/r2dt)^53^. Initial ribosomal RNA (rRNA) models were predicted by AlphaFold3^48^, and initial protein models were built using CryoAtom^54^. For the two non-ribosomal proteins, ARC-P and ARC-A, the atomic models were recognized and constructed using CryoAtom “refine and identify the protein” function combined with NCBI sequence alignment. In brief, the unknown protein backbone was constructed using Ala in Coot v0.9.8.1^55^, and then based on the density of side chain to retrieve in the database and refine based on the sequence of targeted protein. Finally, the model was inspected for completeness and manually built-in regions where the sequences were missing or incorrect. Final models were further subjected to refinement with Phenix.real_space_refine v1.21.2^56,57^. The figures were prepared with ChimeraX v1.8^58^ and compiled using Adobe Illustrator.

### Recombinant protein expression and purification

Full-length coding sequences of ARC-P (GenBank: WYM85633.1), ARC-A (GenBank: WYM85634.1) and eS19 (GenBank: WYM83892.1) were PCR-amplified from *Methanolobus* genomic DNA. ARC-P and ARC-A were cloned into a plasmid containing a C-terminal 6×His-FLAG tag, while eS19 was cloned into pET-507a vectors. Cells were grown in LB medium at 37 °C until OD_600_ reached 0.6–0.8, at which point expression was induced by adding 0.4 mM Isopropyl β-D-1-thiogalactopyranoside (IPTG), and the culture was continued for an additional 3 h at 37 °C. Cells were harvested and resuspended in their respective buffers: ARC-P and ARC-A in lysis buffer A (30 mM HEPES-KOH pH 7.5, 500 mM KCl, 20 mM Imidazole, 1 mM TCEP), and eS19 in lysis buffer B (50 mM HEPES-KOH pH 7.5, 100 mM NaCl, 20 mM Imidazole, 1 mM TCEP). Cells were lysed using a high-pressure homogenizer (1290 bar, 4 passes). After cell lysis and centrifugation, the supernatant was loaded onto a pre-equilibrated Ni-NTA affinity chromatography column. The column was washed with lysis buffer containing 40 mM Imidazole, and the target protein was finally eluted with lysis buffer containing 300 mM Imidazole. Eluted proteins were buffer-exchanged into storage buffers (ARC-P/ARC-A: 30 mM HEPES-KOH pH 7.5, 500 mM KCl, 20% Glycerol. eS19: 50 mM HEPES-KOH pH 7.5, 500 mM NaCl) using desalting columns. Finally, proteins were concentrated, flash-frozen in liquid nitrogen, and stored at -80 °C.

### *In vitro* transcription assays

mRNA for the reporter gene (Nanoluciferase, Nluc) was prepared by PCR amplification of the DNA template, which included a T7 promoter sequence upstream of the Nluc coding sequence. *In vitro* transcription reactions were performed as described previously^59^. Briefly, in a buffer system containing T7 RNA polymerase, nucleoside triphosphates (NTPs), and an RNase inhibitor, incubated at 37 °C for 2 h. The template DNA was removed by DNase I treatment, and the transcribed mRNA was subsequently purified by gel extraction for use in the translation assay. mRNA transcripts were assessed for quality and size by gel electrophoresis, quantified spectrophotometrically (Implen spectrophotometer, Nanophotometer NP80), diluted to 1 μg/µL, aliquoted, and preserved at -80 °C prior to *in vitro* translation reactions.

### *In vitro* translation assays

Recombinantly purified of ARC-P and ARC-A were tested for inhibitory activity in a cell-free translation system. *Methanolobus* cell lysate was prepared using a protocol reported with modifications^50^. Briefly, cells were lysed using a high-pressure homogenizer, and the supernatant was clarified by centrifugation. The 10 μL *in vitro* translation reaction mixtures contained: 5 μL cell lysate, 300 ng Nluc mRNA, 0.1 mM spermidine, 100 µM amino acid mixture, 62.5 μg/mL creatine kinase (CrK), 80 mM creatine phosphate (CrP), 10 mM Mg(OAc)_2_, 100 mM KAC, and 1 μL of the recombinant protein or storage buffer. To determine the role of ATP, 2 mM ATP or solution buffer was added to the IVT system to verify its effect on inhibition of ARC. Reactions were equilibrated at 28 °C for 1 h prior to luciferase activity measurement using a microplate luminometer (BERTHOLD, Centro XS3 LB 960). Lysates and mRNA aliquots from identical preparation lots were used for comparative assays within individual experimental series to control batch variability.

### Ribosome binding assays

Empty ribosomes were obtained by pelleting *Methanolobus* cell lysate through a 0.8 M high-salt sucrose buffer (20 mM HEPES-KOH pH 7.5, 10 mM Mg(OAc)_2_, 1 M KOAC, 1 mM DTT). 10 μM of purified recombinant ARC-P and ARC-A proteins (6×His-FLAG tagged) were incubated with the empty ribosomes at 4 °C for complex formation. The reaction mixture was pelleted through a 0.8 M sucrose cushion (Beckman Coulter MLA130 rotor, 603,000 *g* for 50 min at 4 °C) to isolate ribosomes and associated factors. The ribosome pellet was resuspended in resuspension buffer. The ribosome pellet was then resuspended in resuspension buffer (20 mM HEPES-KOH pH 7.5, 10 mM Mg(OAc)_2_, 30 mM KCl, 1 mM DTT) and used for Western Blot (WB) detection analysis using anti-FLAG tag antibodies (Sigma, A8592-1MG) to confirm co-sedimentation of ARC-P and ARC-A with the purified ribosomes.

### Constitutive and nucleotide-dependent ARC-ribosome association

To investigate the association dynamics of ARC across different physiological states, we performed a series of immunoblotting assays. First, to assess ARC occupancy throughout the life cycle of *Methanolobus*, crude ribosomes were pelleted from cell lysates harvested at distinct growth stages: the logarithmic phase (OD_600_ = 0.2, 0.4, and 0.6), the stationary phase (OD_600_ = 0.9, sampled 12 h after reaching this density), and the late stationary phase (OD_600_ = 0.9, sampled 48 h after reaching this density) by centrifugation through a 0.8 M sucrose cushion. For subunit-specific localization, clarified lysates were fractionated on a 10%–40% (w/v) sucrose gradient at 247,600 *g* for 2.5 h (SW41 rotor). Fractions corresponding to the small ribosomal subunit (SSU) peak were collected and concentrated by ultracentrifugation without an additional sucrose pad. Samples were resolved by 15% SDS-PAGE and transferred to membranes for detection using custom-made polyclonal antibodies specifically targeting ARC-A and ARC-P.

To further evaluate the sensitivity of the ARC-ribosome interface to cellular energy charge, we performed nucleotide-dependent recruitment assays. Fractions corresponding to the 70S peak were obtained via sucrose gradient centrifugation and pelleted via sucrose cushion. The resulting ribosomal pellets were resuspended in grid buffer supplemented with 2 mM adenylates (AMP or ATP) and incubated for 30 min at 15 °C. Ribosomes were subsequently isolated via a 0.8 M sucrose cushion at 603,000 *g* for 50 min (MLA130 rotor) under different concentrations of salt (100mM and 500mM). Pelleted ribosomes were resolved by 15% SDS-PAGE and transferred to membranes for detection using custom-made polyclonal antibodies specifically targeting ARC-A and ARC-P.

### Reconstitution of the AMP-bound 70S-ARC complex

To resolve the structure of the energy-sensing state, purified 70S ribosomes were harvested from the second peak of a 10%–40% sucrose gradient and subjected to a high-salt wash (0.8 M sucrose cushion containing 1 M KOAc) to remove endogenous factors. For reconstitution, ribosomes (0.84 µM) were incubated with recombinant ARC complex (10 µM, pre-incubated with a 100-fold molar excess of AMP) on ice for 15 min. The supernatant was then pelleted through a 0.8 M sucrose cushion. The final pellet was resuspended and supplemented with an additional 50 µM AMP and 0.5 µM ARC complex immediately prior to plunge-freezing on cryo-EM grids. Cryo-EM data collection and subsequent image processing were performed as described in the preceding sections.

### Molecular dynamics simulation

To evaluate the impact of ligand binding on the structural dynamics of the ARC-A/P complex, all-atom molecular dynamics (MD) simulations were performed for three distinct systems: (i) the apo ARC-A/P complex, (ii) the complex bound with three AMP molecules (ARC-A/P-AMP), and (iii) the complex bound with three ATP molecules (ARC-A/P-ATP). The initial structure for the ARC-A/P-AMP system was obtained from a cryo-electron microscopy (cryo-EM) structure (PDB: 21NN), which reveals one AMP molecules bound to the ARC-A and two to the ARC-P. For the ARC-A/P-ATP system, the binding poses of three ATP molecules were predicted using AlphaFold3^48^, with the experimentally resolved AMP sites as spatial references. The apo system was prepared by removing all ligands from the cryo-EM structure.

Prior to simulation, protein structures were prepared using the Protein Preparation Wizard (Schrödinger Suite) to add hydrogen atoms and optimize protonation states at pH 7.4. Ligands (AMP and ATP) were prepared with LigPrep to generate low-energy conformations with appropriate ionization states. Each system was solvated in an orthorhombic box of SPC water molecules, extending 10 Å beyond the protein surface. Systems were neutralized with counter-ions and brought to a physiological salt concentration of 0.15 M NaCl. The OPLS4 force field was applied to all atoms. Energy minimizatin and system equilibration followed the standard Desmond protocol. Subsequently, production MD simulations were conducted for 100 ns per system in the isothermal-isobaric (NPT) ensemble at 300 K and 1.01325 bar. Trajectories were saved at 100-ps intervals. The conformational stability of each system was assessed by calculating the root-mean-square deviation (RMSD) and root-mean-square fluctuation (RMSF) relative to the respective starting structure.

### Construction of the ARC deletion mutant in *Methanolobus*

*Methanolobus* and its derivative strains were cultured anaerobically in ZRKC1 medium (modified from basal medium: 1.0 g/L yeast paste, 1.0 g/L peptone, 1.0 g/L NH_4_Cl, 1.0 g/L CH_3_COONa, 1.0 g/L NaHCO_3_, 0.2 g/L MgSO_4_, 1.0 g/L cysteine hydrochloride, 500 µL/L 0.1% resazurin solution, 1.0 L filtered seawater (pH 7.0) supplemented with 247 mM methanol) at 37 °C under a headspace of N_2_/CO_2_ (80:20, v/v). Solid medium was prepared by adding 1.5% (w/v) agar. When required, puromycin (2.5 or 1.25 µg/mL) was used for selection of genetic transformants.

Gene deletions in *Methanolobus* were generated using a suicide-vector system developed in this study. Approximately 600–800 bp upstream and downstream regions of the target genes, *arcA* and *arcP* (Gene ID: J1 GM001137 and J1 GM001138 in *Methanolobus*) were PCR-amplified from *Methanolobus* genomic DNA to serve as homology arms. A puromycin-resistance cassette was amplified from plasmid pMEV4^60,61^. The three fragments were designed with overlapping termini and assembled into the pMD19-T vector (TAKARA) using Gibson Assembly (NEBuilder HiFi DNA Assembly Master Mix, NEB). The assembled plasmids of pTZC1 (table. S4) were first transformed into *E. coli* DH5α for propagation and verification.

The resulting suicide plasmids (table. S4) were then transformed into *Methanolobus* by polyethylene glycol (PEG)–mediated transformation (described below). Double-crossover homologous recombination events yielded the desired deletion mutants, which were subsquently selected on ZRKC1 media supplemented with puromycin for 7-8 successive sugenerations to ensure complete allelic replacement across all chromosomal copies. Successful gene deletions were confirmed by colony PCR and Sanger sequencing using primers listed in table. S4. For rapid genomic DNA preparation, single colonies were inoculated into liquid medium and grown to late-log phase. One milliliter of cell culture was centrifuged at 13,400 *g* for 1 min, and the pellet was resuspended in 100 μL of sterile distilled water. The suspension was vigorously mixed by pipetting, boiled for 5 min, and the resulting crude lysate was directly used as the PCR template. PCR amplification was performed using 2× Phanta Master Mix (Vazyme, P520), and the amplicons were analyzed by agarose gel electrophoresis and Sanger sequencing to confirm the desired genomic deletions. The growth of wild-type and mutant strains under different conditions was monitored spectrophotometrically by measuring the optical density at 600 nm (OD_600_).

### PEG-mediated transformation and selection of mutant strains

A modified PEG-mediated protocol for transformation of *Methanolobus* was first established in this study. Cells of *Methanolobus* were cultured anaerobically to mid-logarithmic phase (OD_600_ ≈ 0.6), and 5 mL of culture was harvested by centrifugation at 3,000 *g* for 10 min at room temperature. Pellets were gently washed once with 5 mL of oxygen-free transformation buffer (TB; 50 mM Tris base, 350 mM sucrose, 380 mM NaCl, 1 mM MgCl_2_, 1 mM cysteine-HCl, 1 mM DTT, and 0.1 mg/L resazurin, pH 7.5). Cells were resuspended in 380 µL of plasmid DNA solution (2 µg DNA in TB) and mixed with 200 µL of PEG buffer (40% [w/v] PEG 8000 in TB). Following incubation at 37 °C for 1 h, 5 mL of fresh ZRKC1 medium was added to remove residual PEG, and cells were pelleted at 4,000 *g* for 20 min. The pellet was resuspended in 5 mL of fresh ZRKC1 medium and recovered anaerobically at 37 °C for approximately 32 h. Transformants were enriched by serial subculturing for several generations in ZRKC1 medium supplemented with puromycin (1.25 µg/mL) until a stable antibiotic-resistant phenotype indicative of complete allelic replacement was obtained. Puromycin-resistant colonies were subsequently isolated on solid ZRKC1 medium (1.5% agar) and verified by PCR using gene-specific primers.

### Bioinformatic and phylogenetic analysis

Homologs of ARC-P and ARC-A were identified in *Methanolobus* using their respective amino acid sequences to query the UniProt^62^ TREMBL database (release 2025_04). Search parameters included identities >20%, a score >200 bits, and an e-value cut-off of 1 × 10^−13^. All homology search results are presented in Supplementary Tables 4. The identified ARC-P and ARC-A homologs were aligned using MAFFT v7.0^63^. Subsequently, maximum-likelihood phylogenetic trees were constructed with IQ-TREE v2.2.0^64^, employing the best-fit evolutionary models determined by the software and 1,000 ultrafast bootstrap replicates. The resulting trees were visualized using the online tool Interactive Tree of Life (iTOL) v6.0^65^. In addition, the distribution of ARC-P, ARC-A, or both homologs across archaeal lineages was visualized on the GTDB v214 archaeal phylogenetic tree using the iTOL v6.0 web server.

### Conformational dynamics via 3D variability analysis (3DVA)

To resolve the structural heterogeneity and the precise docking transitions of the ARC complex during oxidative stress, we performed 3D variability analysis (3DVA) in cryoSPARC v4.6.2^49^. An initial set of 81,395 particles—comprising vacant and ARC-bound 70S classes from the oxygen-exposed sample—was analyzed using a whole-ribosome mask generated in RELION 4.0.1^52^.

The analysis was performed with a 4 Å low-pass filter, solving for one principal mode to capture the primary conformational movement of the complex. To visualize the transition, 50 frames were generated using the 3D variability display module. The resulting volume series was visualized in ChimeraX v1.8^58^ using the vseries command to characterize the dynamic occupancy and steric remodeling induced by ARC.

## Data availability

The data that support the findings of this study are available from the corresponding authors upon reasonable request. The cryo-EM maps have been deposited in the Electron Microscopy Data Bank and the Protein Data Bank under the following accession numbers: EMD-66877 (stationary phase dataset, *Methanolobus* 70S in non-rotated state with ARC), EMD-66878 (stationary phase dataset, *Methanolobus* 70S in rotated state with P/E-tRNA), EMD-66879 (stationary phase dataset, LSU of the *Methanolobus* ribosome), EMD-66880 (O_2_ stress dataset, *Methanolobus* 70S in non-rotated state with ARC), EMD-66881 (O_2_ stress dataset, *Methanolobus* 70S in non-rotated state with P/P-tRNA), EMD-66882 (O_2_ stress dataset, *Methanolobus* 70S in non-rotated state); EMD-66883 (O_2_ stress dataset, *Methanolobus* 70S in rotated state with A/P P/E-tRNA), EMD-66884 (O_2_ stress dataset, *Methanolobus* 70S in rotated state with P/E-tRNA), EMD-66885 (O_2_ stress dataset, LSU of the *Methanolobus* ribosome), EMD-67845 (AMP dataset, reconstituted *Methanolobus* 70S-ARC complex with AMP). The corresponding atomic coordinates have been deposited in the Protein Data Bank with accession codes 9XHK (*Methanolobus* 70S in non-rotated state with ARC, corresponding map EMD-66877) 9XP4 (*Methanolobus* 70S in rotated state with P/E-tRNA, corresponding map EMD-66878), 9XP5 (LSU of the *Methanolobus* ribosome, corresponding map EMD-66879) and 21NN (reconstituted *Methanolobus* 70S-ARC complex with AMP, corresponding map EMD-67845). Source data are provided with this paper.

## Acknowledgements

The cryo-EM data were collected at the Biomedical Research Center for Structural Analysis of the Shandong University. We thank Amy SY Lee (Harvard University) for her valuable suggestions and critical reading of the manuscript. This work was supported by grants from the National Natural Science Foundation of China (32371351 and 32171291 to W.L., 32393974 to J.L., 42221005 to C.S.), National Key Research and Development Program of China (2025YFF0512900 to J.L., 2022YFA0807100 to W.L.), Nature Science Foundation of Shandong Province (ZR2021QC002 to W.L.) the Shandong Excellent Young Scientists Fund Program (2022HWYQ-025 to W.L.), Taishan Scholars Program (tstp20230637 to C.S., tsqnz20221104 to W.L.), Cutting Edge Development Fund of Advanced Medical Research Institute (GYY2023QY01 to W.L.) and the Cheeloo Youth Program of Shandong University to W.L..

## Contributions

W.L., J.L. and C.S. conceived the study. R.Z. provided methanogenic archaea, performed cultivation, and conducted the phylogenetic analysis. K.W. and X.L. performed the biochemistry experiments including the *in vitro* translation, ribosome binding assay, sucrose gradients and cryo-EM samples preparations with contributions from Q.W. and Y.B.. L.Z. performed the cryo-EM data collection, image processing, model building, and structural analysis with help from C.Y.. L.Z. also performed the CryoDRGN analysis. X.H. performed the molecular dynamics simulation. Y.L. performed genetic editing of *Methanolobus* sp. ZRKC1. L.Z., R.Z., K.W., Y.L. and C.Y. prepared the figures with help from W.L., J.L. and C.S.. W.L., J.L. and C.S. supervised the project and wrote the initial draft of the manuscript. All authors analyzed the data and edited the paper.

