## Supplemental figures and tables for "A dual-factor complex governs archaeal ribosome hibernation by sensing energy status"

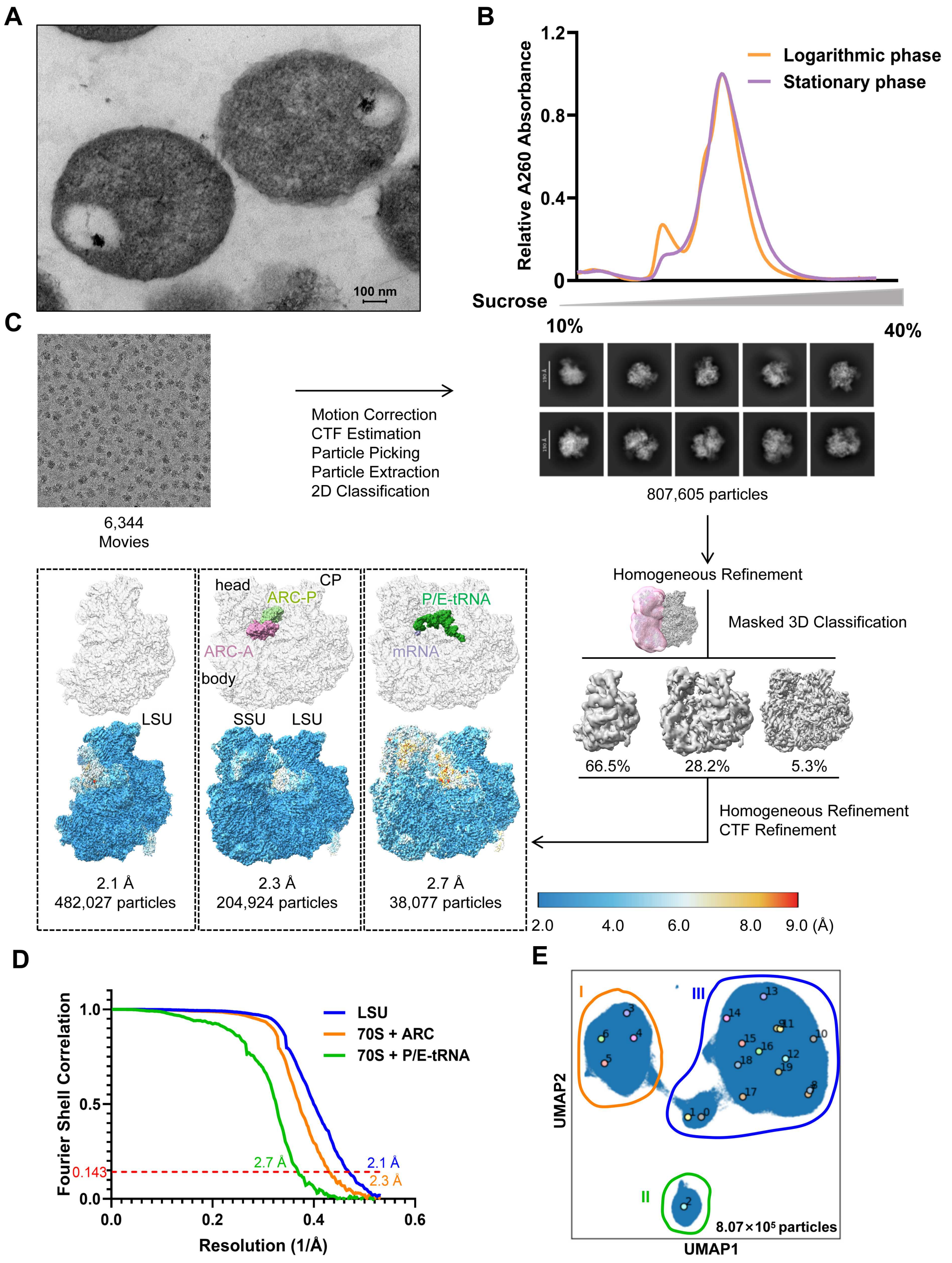
A
100 nm
B
Relative A260 Absorbance
1.2
0.8
0.4
0
Logarithmic phase
Stationary phase
Sucrose
10%
40%
C
6,344
Movies
Motion Correction
CTF Estimation
Particle Picking
Particle Extraction
2D Classification
807,605 particles
Homogeneous Refinement
Masked 3D Classification
66.5%
28.2%
5.3%
Homogeneous Refinement
CTF Refinement
head
CP
ARC-P
ARC-A
body
P/E-tRNA
mRNA
LSU
SSU
LSU
2.1 Å
482,027 particles
2.3 Å
204,924 particles
2.7 Å
38,077 particles
2.0
4.0
6.0
8.0
9.0 (Å)
D
Fourier Shell Correlation
1.0
0.5
0.143
0.0
LSU
70S + ARC
70S + P/E-tRNA
2.7 Å
2.1 Å
2.3 Å
0.0
0.2
0.4
0.6
Resolution (1/Å)
E
I
II
III
UMAP2
UMAP1
8.07×10⁵ particles

**Fig. S1.** Characterization of the *Methanolobus* ribosome and cryo-EM data processing workflow. (**A**) Transmission electron micrograph (TEM) of *Methanolobus* cells. Scale bar, 100 nm. (**B**) Representive sucrose gradient profile (10%–40%) of logarithmic-phase (orange) and stationary-phase (purple) cell extracts. (**C**) The cryo-EM data processing workflow for the stationary phase *Methanolobus* ribosome. Key steps and resulting particle numbers for each classification are indicated. (**D**) Fourier Shell Correlation (FSC) plot showing the overall resolution of the determined cryo-EM structures. The resolution is reported based on the FSC = 0.143 criterion. (**E**) Latent space representations of ribosomal particles as UMAP embeddings after CryoDRGN analysis. Classes are depicted in Roman numbers (e.g., Class I, II, III), map volumes are indicated with Arabic numbers (e.g., Map 1, 2, 3). The total particle number is shown.

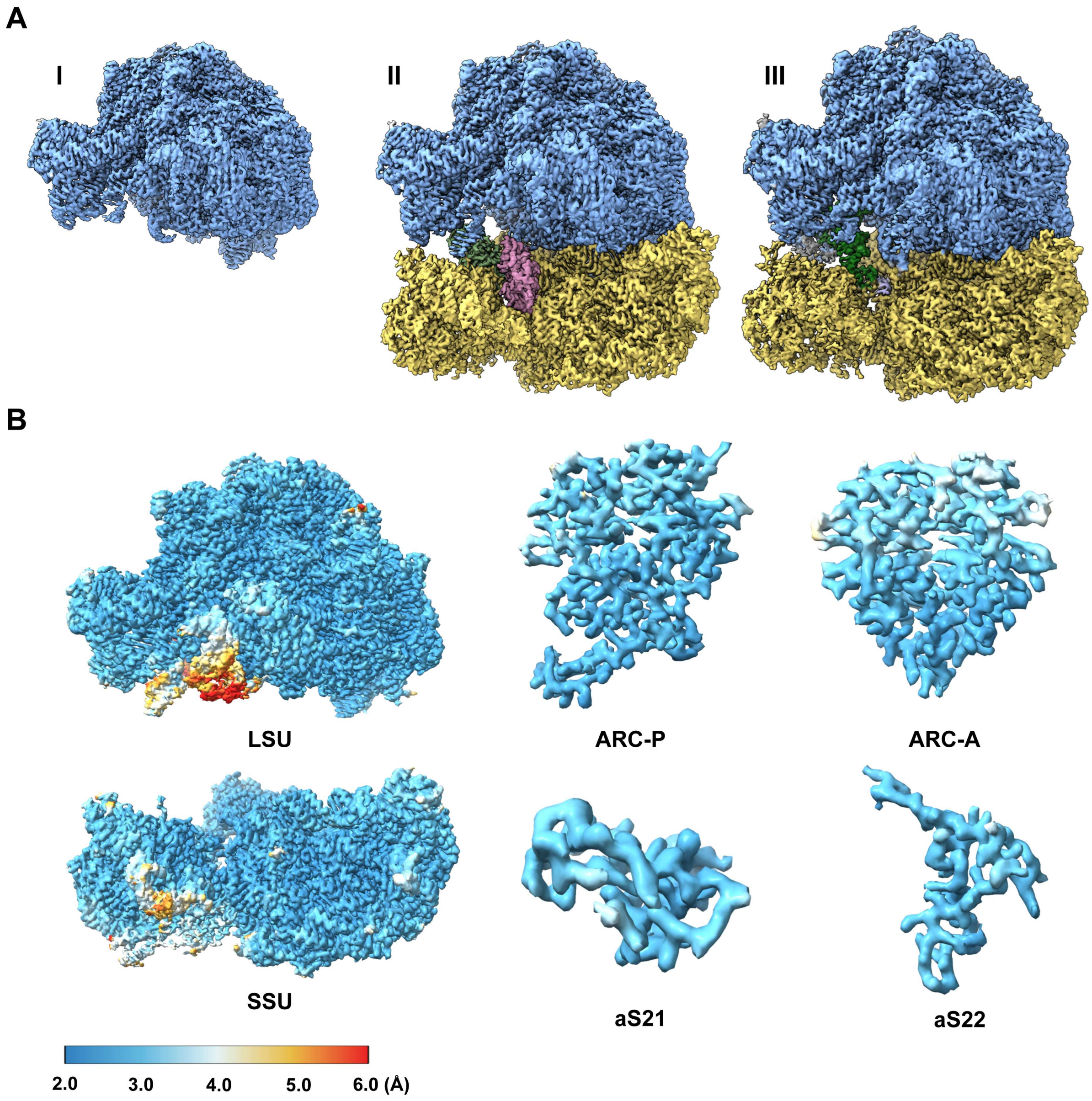


**Fig. S2.** Cryo-EM map quality and detailed features. (**A**) High-resolution cryo-EM maps density of the three main classes of ribosomes identified in stationary phase of *Methanolobus*. (**B**) Local resolution maps of the *Methanolobus* ribosomal components, including the LSU, SSU, ARC-P, ARC-A and aS21, aS22.

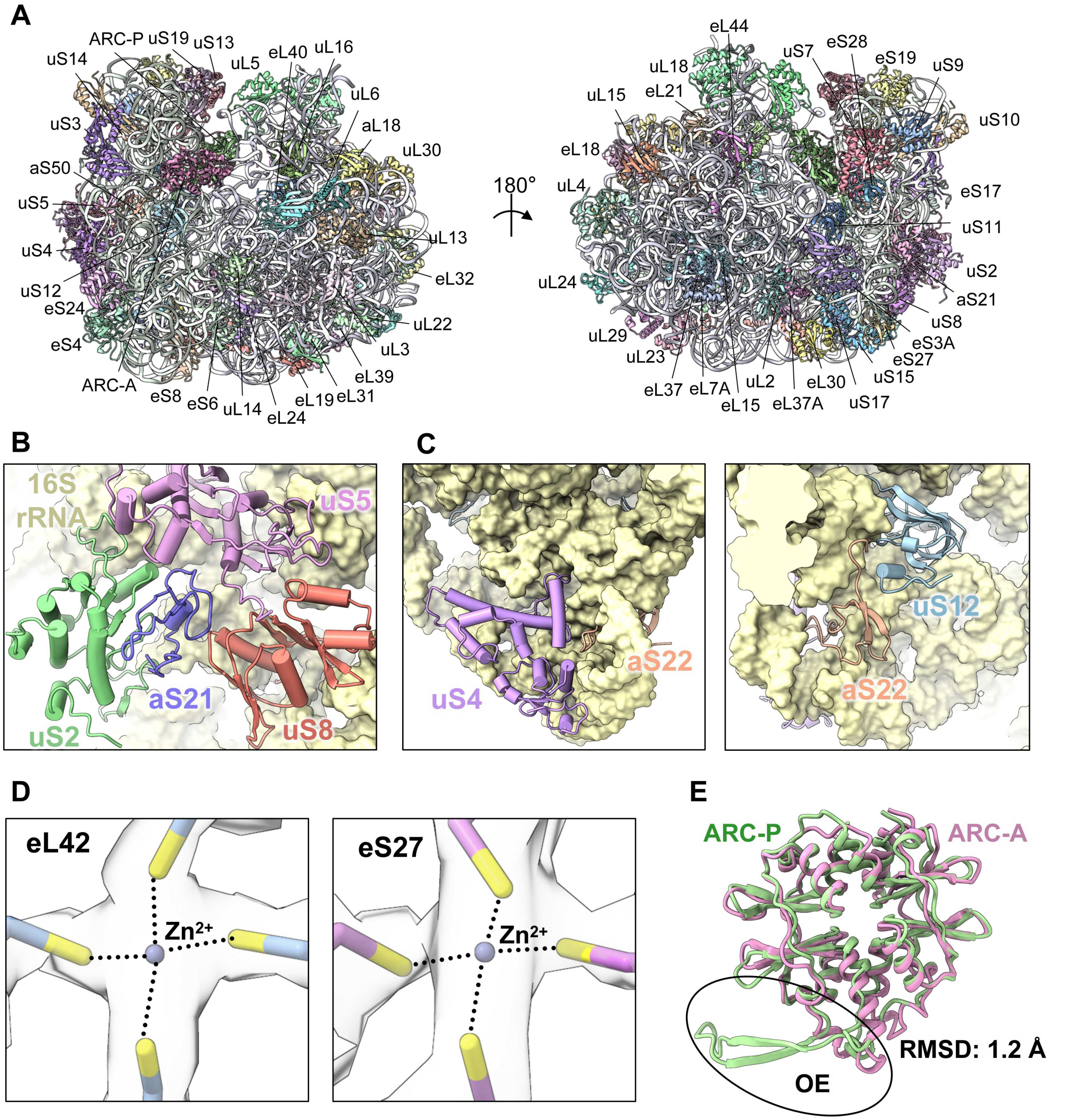


**Fig S3.** Structural characteristics of *Methanolobus* ribosome and ARC. (**A**) Atomic model of the *Methanolobus* 70S ribosome. rRNA and ribosomal proteins are shown in cartoon. (**B**) Localization of the archaea-specific ribosomal protein aS21 (purple) on the SSU. aS21 is located adjacent to ribosomal proteins uS5 (magenta), uS8 (salmon), and uS2 (green). (**C**) Localization of the conserved but previously unannotated ribosomal protein aS22 (wheat) on the SSU. Neighboring ribosomal proteins and rRNA are labeled. (**D**) Detailed views of the coordinated $Zn^{2+}$ ion binding sites, aligned with conserved sites in other archaeal ribosomes[30]. (**E**) Structural alignment of ARC. ARC-P is superimposed on ARC-A with calculated Root Mean Square Deviation (RMSD) value indicated.

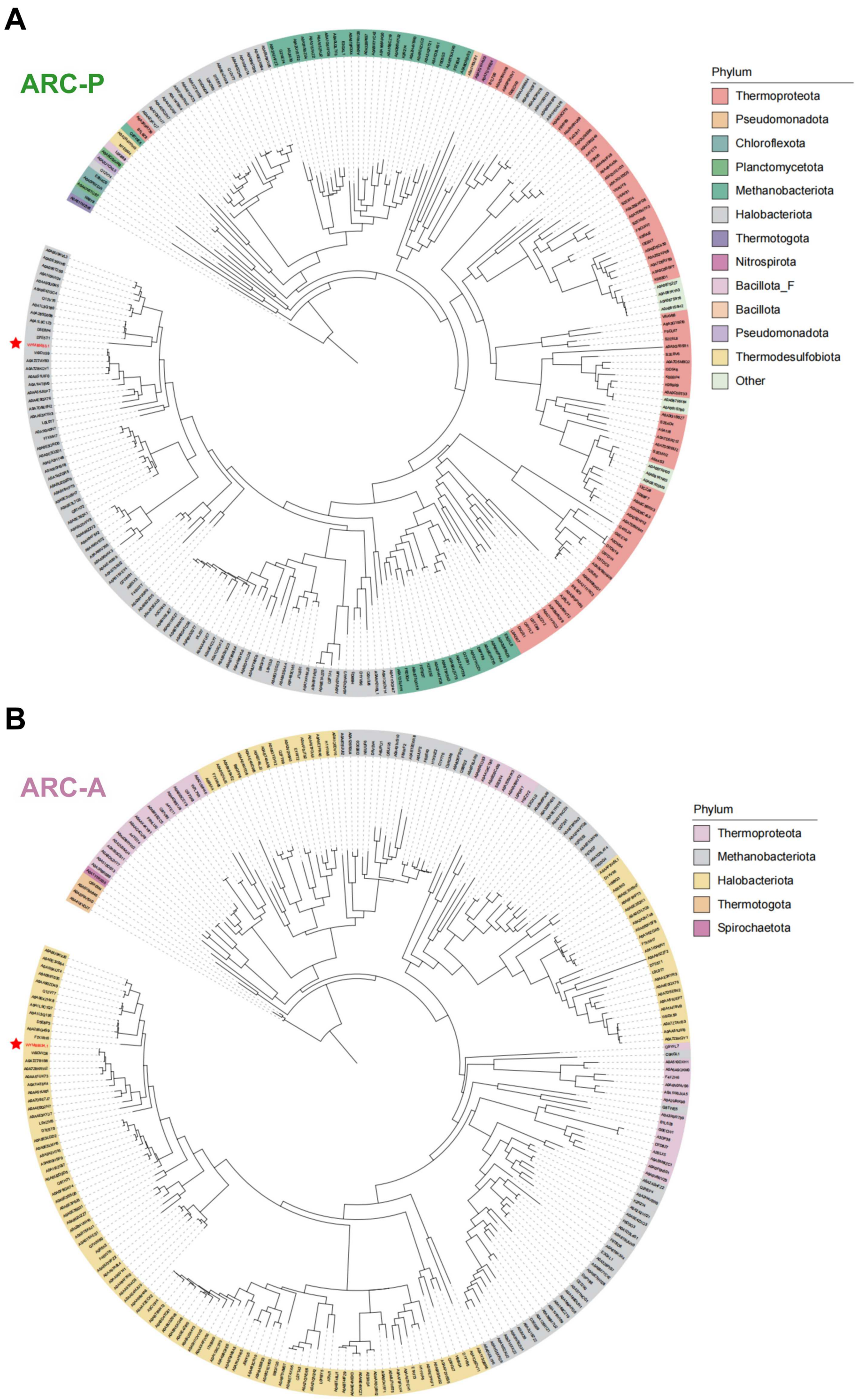


**Fig. S4.** Phylogenetic distribution of ARC-P and ARC-A homologs. (**A**) Phylogenetic distribution of ARC-P homologs across the domain Archaea and Bacteria. (**B**) Phylogenetic distribution of ARC-A homologs across the domain Archaea and Bacteria.

*Methanolobus* is indicated by a red star.

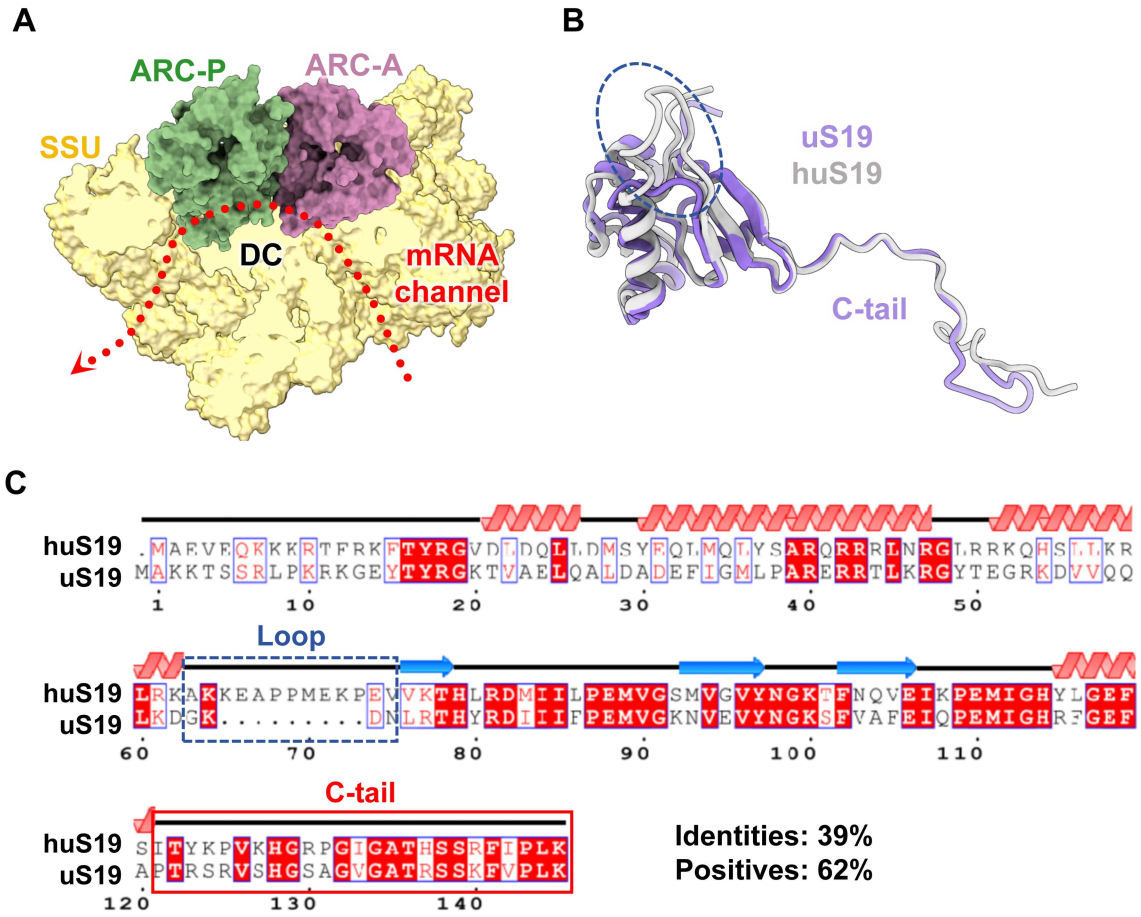


**Fig S5.** Localization of ARC and structural comparison of uS19. (**A**) Overview of ARC anchoring to the mRNA channel and the decoding center. (**B**) Comparison of the tertiary structures of uS19 from *Methanolobus* and human (gray, PDB: 6Y0G[41]). A section of the unstructured loop that is distinctly absent in the *Methanolobus* protein is indicated by a blue dashed box. (**C**) Comparison of the primary and secondary structures of uS19 from *Methanolobus* and human. Black lines highlight the disordered regions, red helices represent α-helix regions, and blue arrows represent β-sheet regions. The unstructured loop absent in *Methanolobus* is indicated by a blue dashed box. The conserved C-tail is indicated by a solid red box.

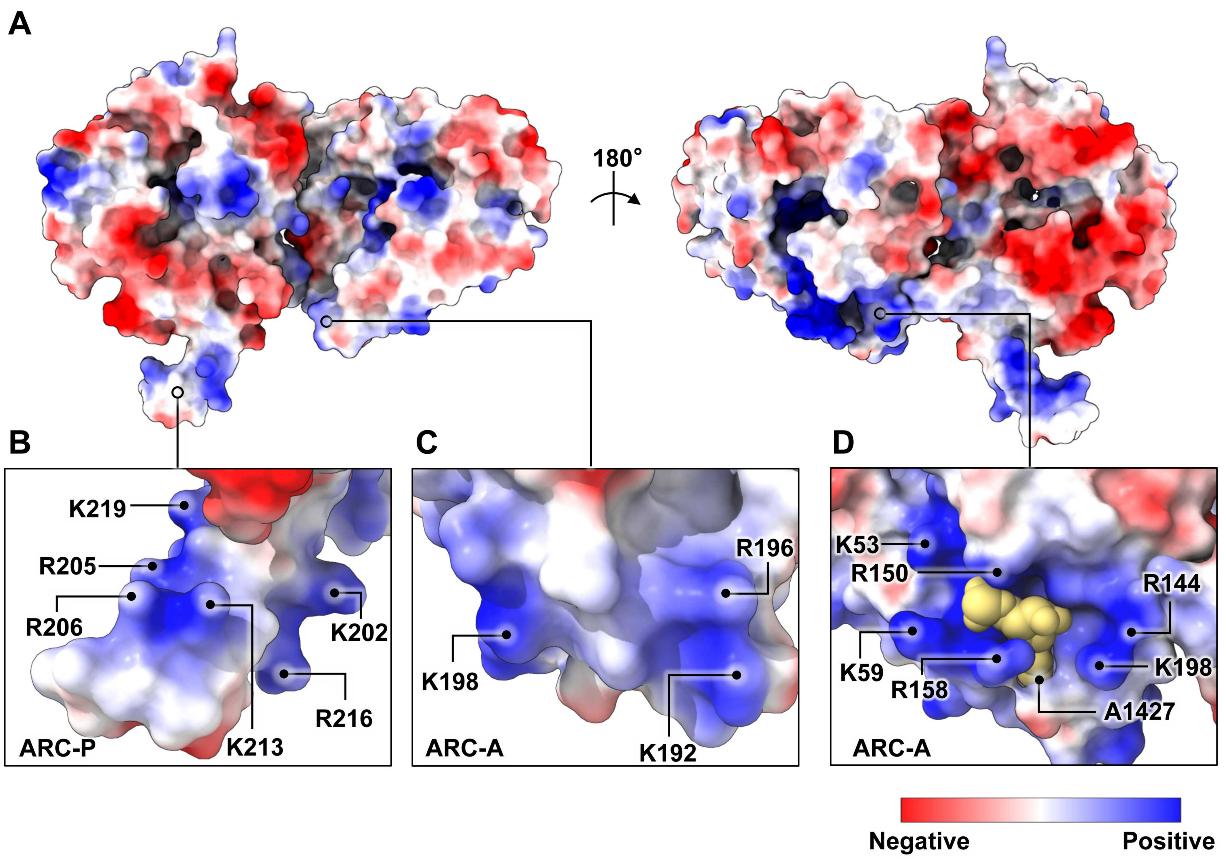


**Fig. S6.** Surface electrostatic potential of ARC. (**A**) Overall electrostatic surface potential of ARC. (**B and D**) Close-up views of the OE of ARC-P (B) and ARC-A (C), showing the highly basic (blue) clusters. (**D**) Electrostatic potential of h44 pocket. Basic residues involved in OE anchoring are indicated; the flipped A1427 base is shown in sphere representation. Potential scale: red (-10 kT/e) to blue (+10 kT/e).

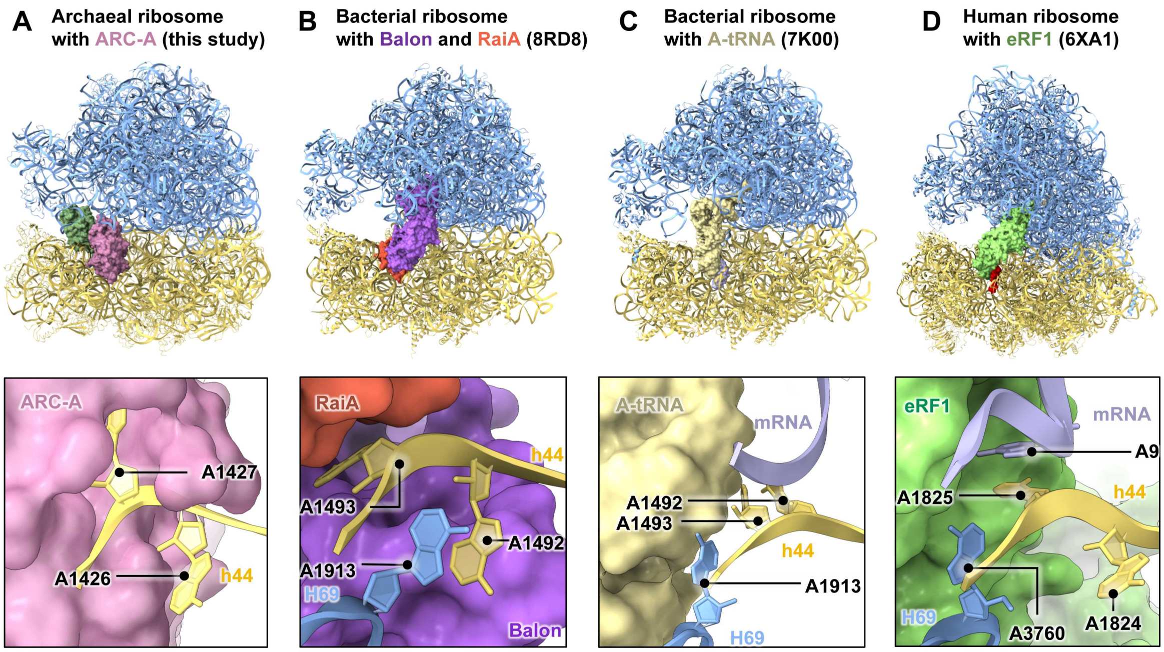


**Fig. S7.** Evolutionary convergence of A-site blockade and h44 remodeling. Structural comparison of A-site occupation mechanisms. (**A to D**) Top panels show the entire ribosome with the bound factor (ARC-A, Balon, A-tRNA and eRF1 in surface). The bottom panels provide zoom-in views of the decoding center (h44), highlighting the conformation of the monitoring base A1427 (A1493 in *E. coli*, A1825 in human). (**A**) ARC-A binding mode. (**B**) Balon binding mode (PDB: 8RD8[10]). (**C**) Canonical A- tRNA binding mode (PDB: 7K00[16]). (**D**) eRF1 binding mode (PDB: 6XA1[44]).

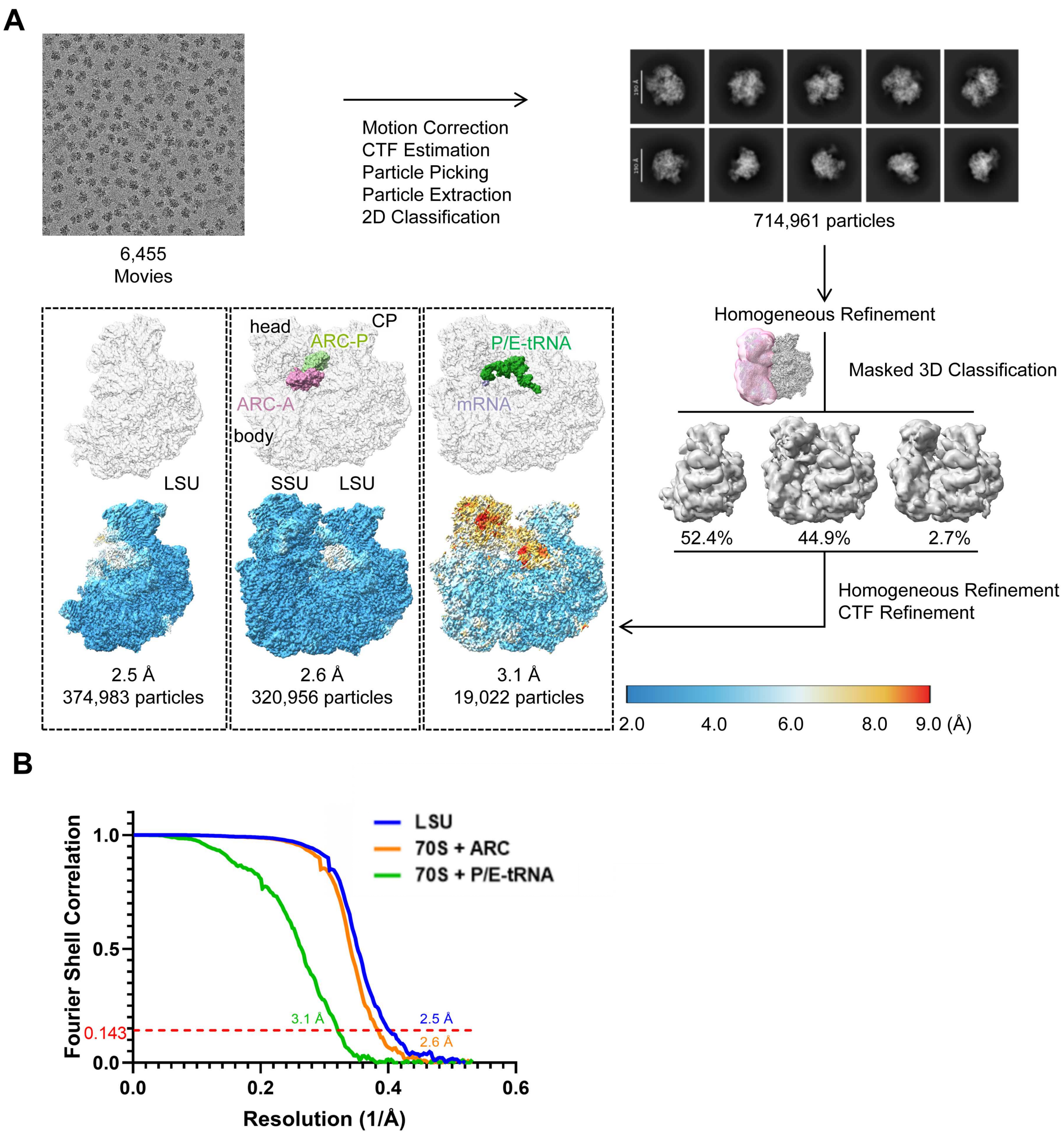


**Fig. S8.** Cryo-EM data processing of the AMP-bound sample. (**A**) Cryo-EM data processing workflow for the *Methanolobus* 70S ribosome incubated with ARC in the presence of 1 mM AMP. (**B**) FSC curve for the refined ribosome structure from the aerobic stress sample. The overall resolution is reported based on the FSC = 0.143 criterion.

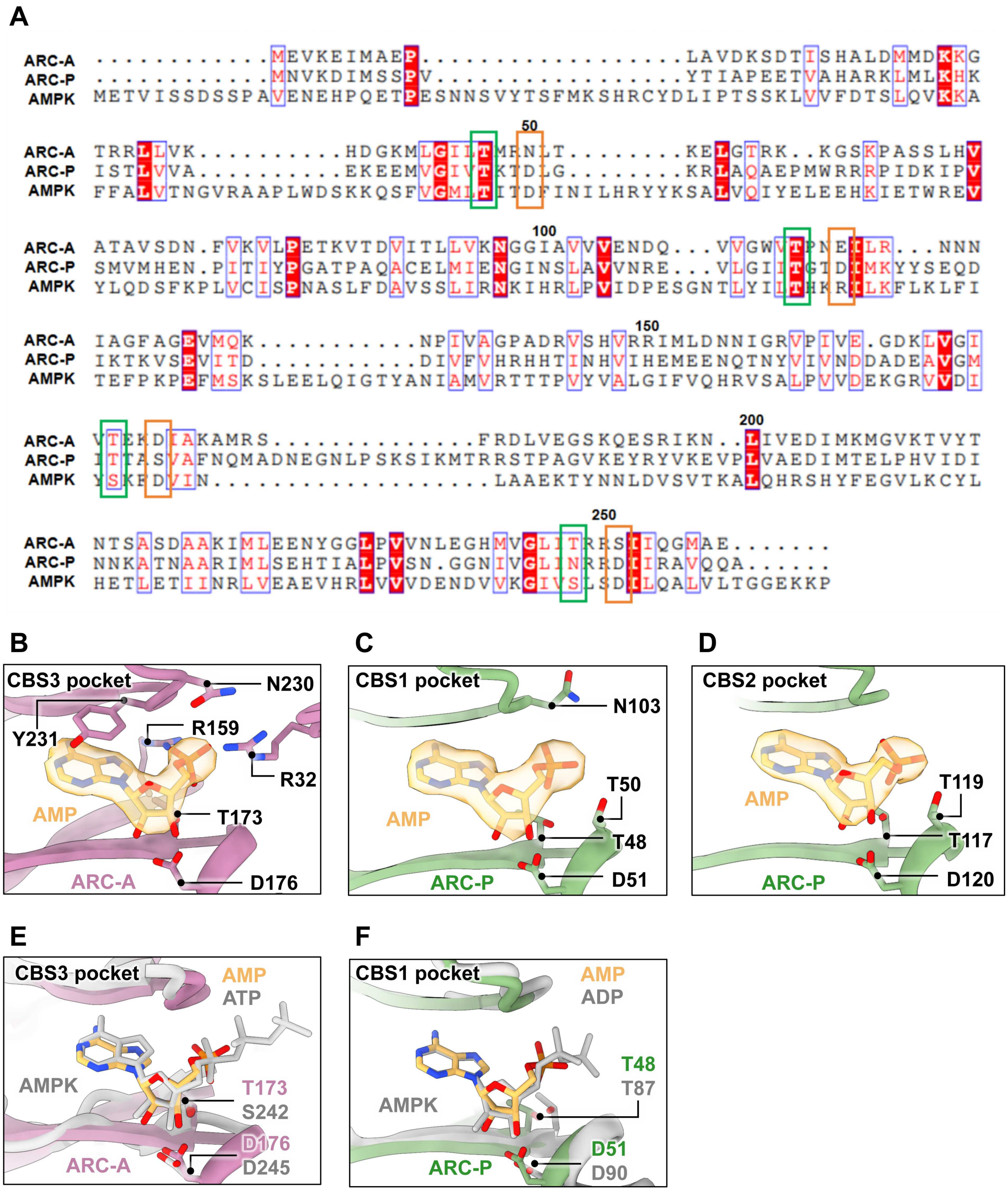


**Fig. S9.** Conservation of CBS pockets and the adenosine phosphate binding. (**A**) Sequence alignment between *Methanolobus* ARC subunits and the human AMPK γ-subunit. The conserved Threonine (T) and Aspartate (D) are indicated by green and orange boxes, respectively. (**B to D**) Detailed views of AMP coordination in ARC-A

CBS3 (B), ARC-P CBS1 (C), and ARC-P CBS2 (D). AMP molecules are shown within their corresponding cryo-EM densities. (**E to F**) Structural superposition of ARC and AMPK (PDB:7JHG) at the conserved CBS pocket. (**E**) Alignment between ARC-A (CBS3) and AMPK (CBS3), showing the relative positioning of bound AMP (ARC-P) and ATP (AMPK). (**F**) Alignment between ARC-P (CBS1) and AMPK (CBS1), showing the relative positioning of bound AMP (ARC-P) and ADP (AMPK).

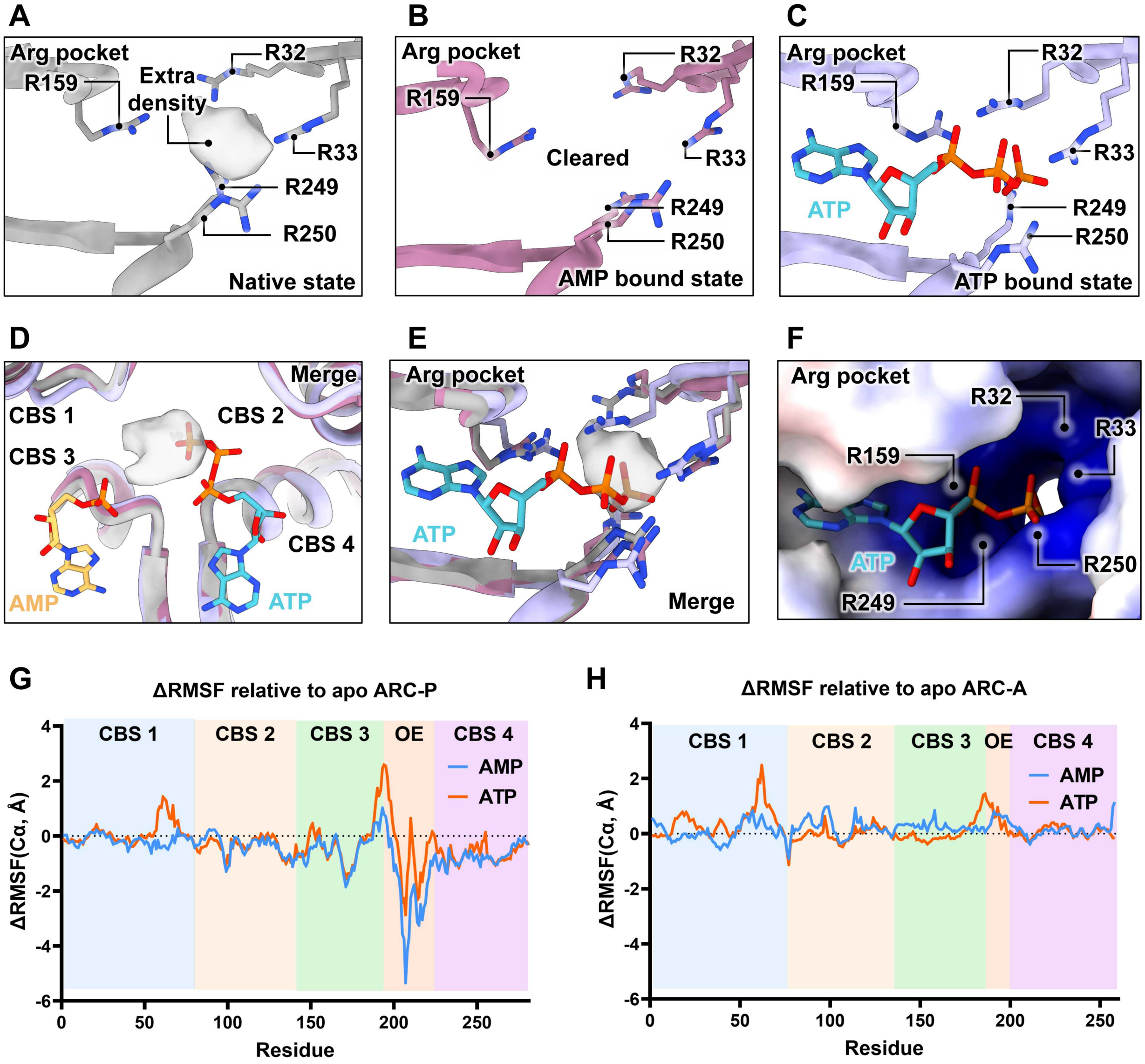


**Fig. S10** Nucleotide-induced conformational dynamics and coordination of the ARC-A sensing pocket. (**A to C**) Zoom-in views of the ARC-A arginine-rich pocket in the native (A), AMP-bound (B) and ATP-bound (C) states. Key arginine residues are shown in sticks. (**D and E**) Structural superimposition of arginine-rich pocket across distinct functional states. (**F**) Structural model of ATP (predicted by AlphaFold 3) docked within the ARC-A sensing site, illustrating the penetration of the polyanionic triphosphate moiety into the basic environment of the arginine pocket. (**G and H**) The Root Mean Square Fluctuation (RMSF) analysis from MD simulations, illustrating the ligand-dependent dynamics of ARC relative to the apo state.

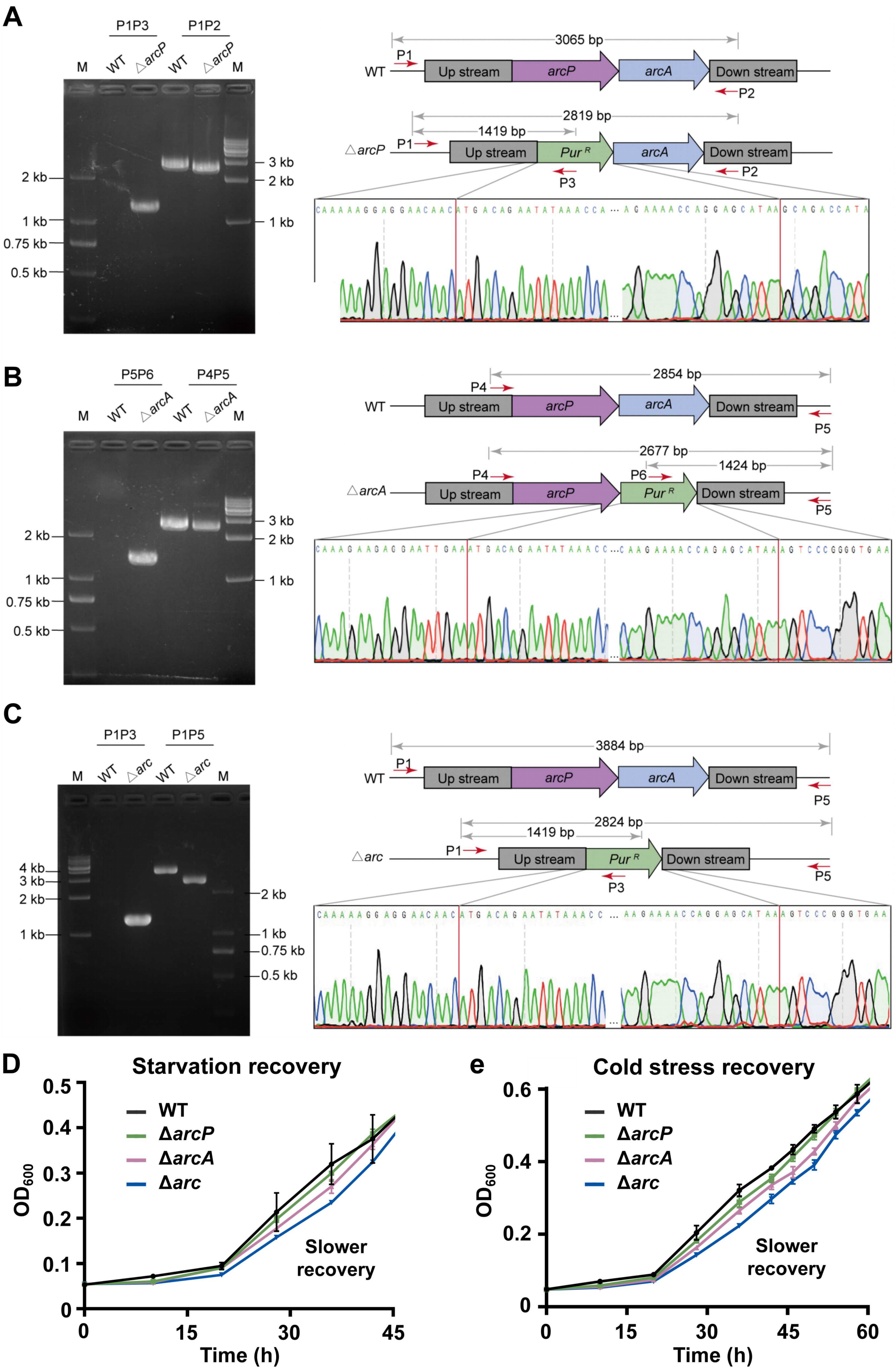
A
P1P3
P1P2
M
WT
△arcP
2 kb
1 kb
0.75 kb
0.5 kb
3 kb
3065 bp
P1
Up stream
arcP
arcA
Down stream
P2
2819 bp
1419 bp
Pur R
P3
B
P5P6
P4P5
△arcA
2854 bp
P4
P5
2677 bp
P6
1424 bp
C
P1P5
△arc
4 kb
3884 bp
2824 bp
D
Starvation recovery
WT
ΔarcP
ΔarcA
Δarc
OD600
Time (h)
Slower recovery
e
Cold stress recovery

**Fig. S11.** Construction and growth analysis of the *arc*-deficient strains. (**A to C**) PCR and sequencing verification of the Δ*arcP* (A)*,* Δ*arcA* (B) *and* Δ*arc* (C) mutant strains obtained from puromycin-resistant transformants. (**D and E**) Zoom in views of the growth curves focusing on the early recovery phase of WT and mutant strains following starvation recovery (D) and cold stress recovery (E).

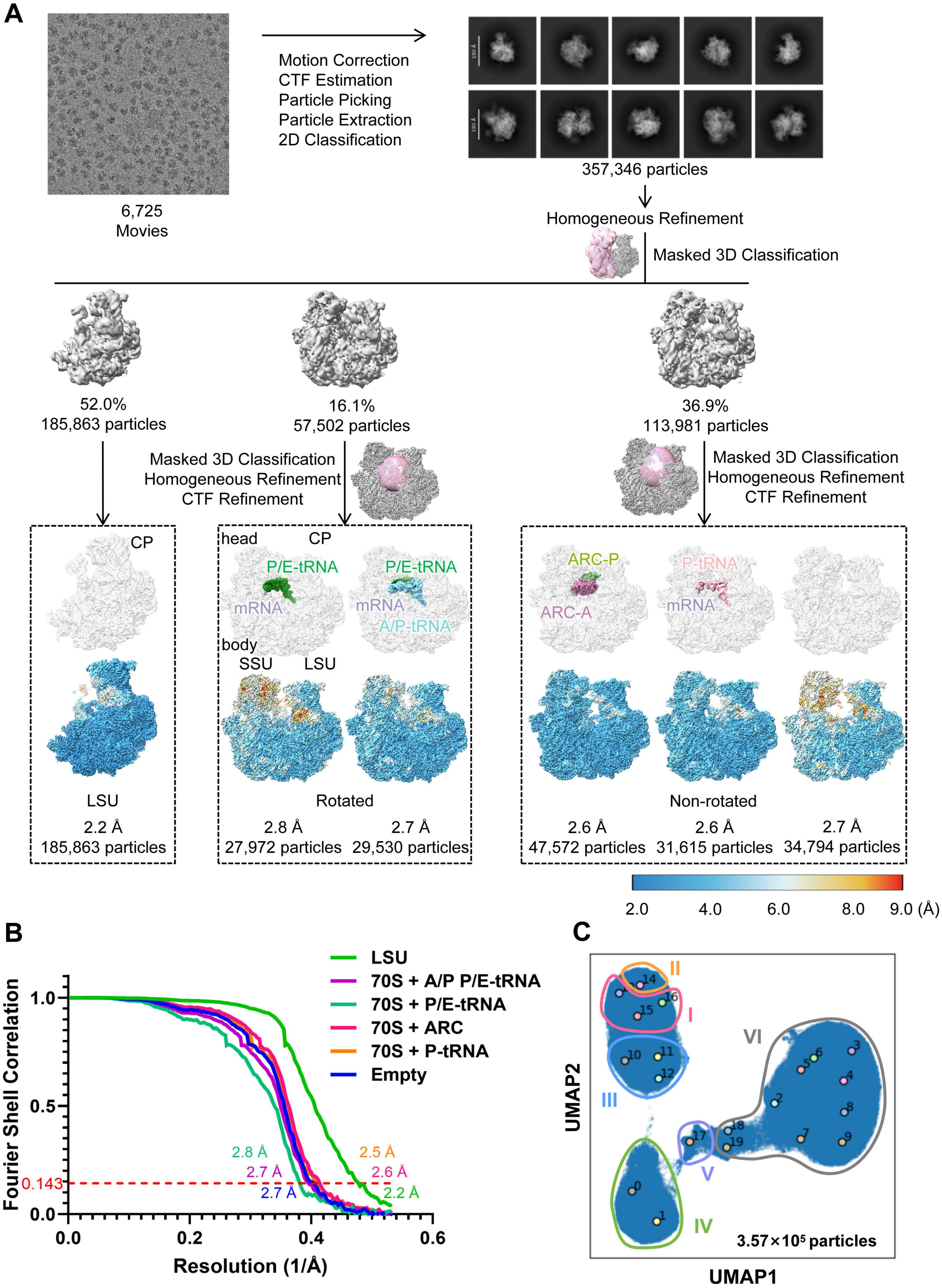
A
6,725
Movies
Motion Correction
CTF Estimation
Particle Picking
Particle Extraction
2D Classification
357,346 particles
Homogeneous Refinement
Masked 3D Classification
52.0%
185,863 particles
16.1%
57,502 particles
36.9%
113,981 particles
Masked 3D Classification
Homogeneous Refinement
CTF Refinement
Masked 3D Classification
Homogeneous Refinement
CTF Refinement
CP
head
CP
P/E-tRNA
P/E-tRNA
mRNA
mRNA
A/P-tRNA
body
SSU
LSU
ARC-P
ARC-A
P-tRNA
mRNA
LSU
2.2 Å
185,863 particles
Rotated
2.8 Å
27,972 particles
2.7 Å
29,530 particles
Non-rotated
2.6 Å
47,572 particles
2.6 Å
31,615 particles
2.7 Å
34,794 particles
2.0
4.0
6.0
8.0
9.0 (Å)
B
Fourier Shell Correlation
Resolution (1/Å)
LSU
70S + A/P P/E-tRNA
70S + P/E-tRNA
70S + ARC
70S + P-tRNA
Empty
0.143
2.8 Å
2.7 Å
2.7 Å
2.5 Å
2.6 Å
2.2 Å
C
UMAP2
UMAP1
I
II
III
IV
V
VI
3.57×10⁵ particles

**Fig. S12.** Cryo-EM data processing and analysis of the acute oxygen stress sample. (**A**) Data processing workflow for the 70S ribosome isolated from the culture subjected to oxygen exposure. (**B**) FSC curve for the refined ribosome structure. The overall resolution is reported based on the FSC = 0.143 criterion. (**C**) CryoDRGN analysis of the sample heterogeneity, with UMAP embeddings showing the distribution of ribosomal classes.
